# Targeted Airborne eDNA Detection of Pest Wallabies: Effects of Sampler Type and Distance

**DOI:** 10.1101/2025.10.26.684691

**Authors:** Gracie C Kroos, Kristen Fernandes, Philip John Seddon, Travis Ashcroft, Neil Gemmell

## Abstract

Bennett’s wallabies *Notamacropus rufogriseus*, introduced to New Zealand from Australia in the late 1800s, strongly exemplify the detection challenges posed by invasive terrestrial species that are rare, cryptic, and highly mobile. Across their invasive range, *N. rufogriseus* occupy large landscapes at low densities, making their surveillance challenging. Recent research has demonstrated that airborne environmental DNA (eDNA) can rapidly identify terrestrial vertebrate diversity in an area. Leveraging these findings, we investigate the utility of airborne eDNA for the targeted monitoring of *N. rufogriseus*, using a novel, probe-based quantitative PCR assay. The effects of filtration material, collection method (active versus passive), distance from the source, and environmental conditions were examined for their effects on airborne detection probability, using a captive population of wallabies in a controlled park setting. A total of 110 airborne samples were collected, 55 with active (battery-powered fan) samplers and 55 passive (non-powered) samplers, across six distinct experimental periods at distances of 0, 10, 100, and 1000 metres from the closest known source of wallaby DNA. Filters designed to capture coarse particles (>10 µm) significantly improved detection rates and DNA recovery for actively collected samples, compared to filters targeting finer particles (1–10 µm). Active samplers significantly outperformed passive samplers in overall detection rates, particularly at shorter ranges from the target. Distance from the source had a significant negative effect on detection probability. Detection rates declined sharply beyond 10 metres but remained possible up to 1 kilometre from the source for both collection methods. These findings demonstrate that airborne eDNA can detect terrestrial vertebrate species at ecologically relevant distances, supporting its potential for landscape-scale surveillance. Notably, these results underscore the importance of optimising sampler design when applying airborne eDNA for targeted species monitoring.

## 1 INTRODUCTION

Environmental DNA (eDNA) has significantly advanced the monitoring of elusive and cryptic terrestrial vertebrate species, offering comparable or superior results to traditional survey methods such as camera traps and field sign surveys, while requiring less labour, taxon-specific expertise, and specialised field equipment, enabling broader-scale applications (Ushio et al., 2017; Leempoel et al., 2020; Mena et al., 2021). However, the effectiveness of eDNA is highly dependent on the species, environment, substrate selection and assay sensitivity (Sales et al., 2020; Croose et al., 2023). Targeted quantitative PCR (qPCR) assays typically provide higher sensitivity for detecting low-density taxa and are more cost-effective, supporting larger sample sizes, wider geographic coverage, streamlined protocols, and simpler interpretation (Blackman et al., 2020). Targeted assays are especially meaningful for invasive species management, where non-detection has serious consequences, such as early invasive species detection (Allen et al., 2021; Morisette et al., 2021) and post-eradication monitoring (Carim et al., 2019), whereby eDNA can guide the deployment of conventional surveys (Furlan et al., 2019).

Air presents a powerful medium for eDNA sampling across diverse environments, with growing evidence supporting its broad potential and effectiveness (Serrao et al., 2021; Lynggaard et al., 2022; Garrett et al., 2023; Polling et al. 2024). This approach is particularly valuable for monitoring terrestrial systems, where eDNA tends to be patchily distributed, making the choice of sample type critical for detecting rare species (Van Der Heyde et al., 2020). Airborne eDNA is both scalable and cost-effective, with strong potential to enhance the early identification of invasive species, particularly in habitats with limited water availability or when species ecology is less water-dependent (Roger et al., 2022; Johnson et al., 2023).

While airborne eDNA holds significant promise for monitoring terrestrial species, several key questions remain. A primary concern is the uncertainty surrounding the spatial resolution of airborne detection, since air can travel long distances and traverse complex pathways to reach a site (Lynggaard et al., 2024). Despite growing interest in the field, the processes underlying the deposition, transport, and degradation of airborne eDNA remain largely unknown (Tulloch et al., 2025). However, airborne particulates are known to exhibit spatial and temporal confinement (Craine et al., 2017; Sherwood et al., 2017), much like the localised distribution patterns of eDNA observed in some aquatic systems (Bruce et al., 2021). Airborne eDNA exhibits strong spatial-temporal signals in both controlled zoo enclosures (Clare et al., 2022) and natural settings (Roger et al., 2022; Lynggaard et al., 2022; Lynggaard et al., 2024; Tournayre et al., 2025). Despite promising evidence that transport of airborne eDNA is generally constrained by local topography, with long-distance movement possible but rare, the spatial resolution of targeted airborne eDNA monitoring schemes remains unclear. This uncertainty is further compounded by the fact that the production, persistence, and degradation of airborne eDNA are influenced by environmental variables such as temperature, humidity, rainfall, and wind conditions (Barnes et al., 2014; Shogren et al., 2017; Harrison et al., 2019; Barnes et al., 2021; Johnson et al., 2023).

Moreover, airborne sampler design plays a critical role in detectability. Active air samplers use powered systems such as vacuums or pumps to draw air through filters, allowing for higher collection efficiency through processing large air volumes over short timeframes (Mainelis, 2020). Simple, battery-powered samplers have proven highly effective at collecting airborne eDNA for a wide range of taxa while being low-cost and easy to deploy (Garrett et al., 2023; Lynggaard et al., 2024). However, these systems are often limited by power requirements, particularly in remote or long-term monitoring scenarios (Roger et al., 2022). Whereas, passive samplers rely solely on wind to deliver particles to unpowered collection surfaces, offering a low-cost, low-maintenance option ideal for long-term or remote use (Johnson et al., 2021, 2023). However, detection efficiency may be lower for short-term deployment, as well as more variable, depending on environmental factors like wind speed (Mendez et al., 2016). Thus, there is a need to directly compare passive and active collection methods, balancing logistical advantages against differences in air volume processing (Johnson et al., 2019, 2023). Additionally, the efficiency of eDNA capture is influenced by filter material and pore size (Bruce et al., 2021). Further research is needed to understand how filter construction impacts detection sensitivity and DNA recovery in airborne applications, which are likely to vary by species, making it an important consideration for targeted, species-specific assays.

Bennett’s wallabies *Notamacropus rufogriseus* (Desmarest, 1817) were introduced to New Zealand in the 1870s for sport and commercial purposes (Latham et al., 2021). Since their introduction, *N. rufogriseus* have thrived in the absence of natural predators, facilitating their rapid spread into neighbouring areas, which has necessitated acute control measures (Latham, Latham & Warburton, 2019; Latham et al., 2021). Wallabies exert significant impacts on primary production in their invasive range, particularly on farming and forestry, as well as impacting the composition of native plant communities by selective browsing of seedlings (Latham et al., 2020). To the detriment of wallaby control programmes, which currently aim to suppress populations within designated containment areas, significant challenges lie in applying operational resources to detect and eliminate the wallabies known to persist outside these containment areas (Tipu Mātoro National Research Plan for dama, parma and Bennett’s wallabies in New Zealand, 2024). Individuals at range boundaries exist at low densities and are mobile across extremely large and varied landscapes leading to spatial heterogeneity in monitoring effort (Latham, Latham & Warburton, 2023). Surveillance is further hindered by the cryptic, elusive, mobile, and semi-solitary nature of *N. rufogriseus*, which exhibit crepuscular activity (most active under low light conditions) (Hickling & Day, 2024).

Building on these findings, we evaluate the utility, sensitivity, and accuracy of airborne eDNA for detecting Bennett’s wallaby *N. rufogriseus* in its invasive range in New Zealand. First, we develop a novel, highly sensitive species-specific qPCR assay for *N. rufogriseus* and validate the assay in controlled, open-air environments that realistically simulate natural conditions while allowing precise control over individual numbers and sampling distances from the DNA source. Next, we assess optimal filter materials and collection techniques for capturing airborne eDNA from *N. rufogriseus*. Additionally, we investigate the dispersal distance of wallaby airborne eDNA signals from the nearest known source, alongside the influence of environmental conditions on detectability.

## 2 MATERIALS AND METHODS

### 2.1 Primer Design and Validation

Candidate primers and probes were designed and tested using sequence data from the NCBI GenBank database https://www.ncbi.nlm.nih.gov/genbank/; accession numbers: <u>KY996499</u>, <u>KJ868122</u>, <u>JN003393</u> [*Notamacropus rufogriseus*]. Primers were designed to target the mitochondrial NADH dehydrogenase 2 gene (MT-ND2 gene), chosen for targeted assay development following a scan of the entire mitochondrial genome in Geneious Prime 2023.2.1 (https://www.geneious.com). The MT-ND2 gene revealed sufficient representation in online databases and high levels of inter-specific sequence variation among closely related species *Notamacropus rufogriseus* (Bennett’s wallaby) and *Notamacropus eugenii* (dama wallaby), as observed from additional NCBI GenBank records; accession numbers: <u>KJ868119</u>, <u>JN003390</u>, <u>CM051844</u> [*Notamacropus eugenii*].

Primers and probes were designed in Geneious Prime 2023.2.1 targeting short fragments (80-150 bp) as is recommended for amplifying DNA from environmental samples (De Brauwer et al., 2023). In-built Primer3 software in Geneious Prime and OligoAnalyzer tool (Integrated Gene Technologies) were used to check the annealing temperature (T_m_), GC content, and possible secondary structures. PCR products were aligned against a custom-built reference database comprising all available full-length animal mitochondrial genomes in NCBI within the programme CRABS v0.1.8 (Jeunen et al., 2023). Forward and reverse primer sequences were designed to have 0 mismatches to the target, and at least one mismatch in each of the forward and reverse regions to nontarget species. In addition, the probe sequence was designed to have at least two mismatches to nontarget species (Klymus et al., 2020; De Brauwer et al., 2023). Primer pairs and minor groove binding (MGB) probe deemed suitable by *in silico* analyses were ordered (Integrated Gene Technologies).

Primer specificity was then evaluated under controlled laboratory conditions using qPCR on a QuantStudio 3.0 Real-time PCR instrument (Life Technologies), with genomic DNA extracted from frozen ear tissue of a single *N. rufogriseus* individual collected from mainland New Zealand. Genomic DNA from 11 closely related/co-occurring mammalian species sourced from New Zealand were tested for any non-specific amplification: dama wallaby, *Notamacropus eugenii*; parma wallaby, *Notamacropus parma*; common brushtail possum, *Trichosurus vulpecula;* red deer, *Cervus elaphus;* black rat, *Rattus rattus;* pig, *Sus scrofa;* cow, *Bos taurus;* sheep, *Ovis aries;* house mouse, *Mus musculus;* human, *Homo sapiens;* and European hedgehog, *Erinaceus europaeus*. All qPCR reactions to test assay specificity and sensitivity were set up in a UV hood with exclusive use of filter tips. Technical replicates were run in triplicate for each DNA sample. Each plate also contained no-template control samples (nuclease-free water in place of template DNA) in triplicate. Reactions were carried out in 20 µl volumes; each containing 0.8 µl of the forward and reverse primers, and 0.2 µl of the probe (all 10 µM working stock),10 µl 2x SensiFAST probe mix (ThermoFisher Scientific), 6.2 µl nuclease-free water, and 2 µl template DNA. Gradient PCR was performed to determine the optimal annealing temperatures of each assay. Initial thermocycling conditions were as follows: an initial denaturation at 95 °C for 3 minutes, followed by 40 cycles of denaturation at 95 °C for 15 seconds, annealing at 55°C, 58 °C, and 60 °C for 30 seconds; and an extension at 72 °C for 45 seconds. PCR products were then analysed by agarose gel electrophoresis to assess amplification efficiency and specificity.

The sensitivity of the best-performing assay was tested by calculating the limit of detection (LOD), defined as the lowest standard concentration of target DNA that could be positively detected and quantified in 95% of cases (Bustin et al., 2009) and the limit of quantification (LOQ), defined as the lowest standard concentration of target DNA that could be quantified with a coefficient of variation (CV) value below 35% (Klymus et al., 2020). A standard curve was generated using synthetic, double-stranded oligonucleotides of the *N. rufogriseus* MT-ND2 gene, consisting of the 106 bp PCR product flanked by additional 20 bp sequences on either end (total length 146 bp). This fragment was inserted into a research-grade DNA plasmid (pUC57 vector), which was isolated from *E. coli* strain DH5α, resulting in a total plasmid size of 2,856 bp (Table S1). The plasmid was delivered lyophilised at 10 µg (GenScript), and synthetic target DNA was diluted to a concentration of 10 ng/µl in low EDTA TE buffer (Thermo Fisher Scientific), then further diluted to make a 10-fold dilution series starting at 64880 copies – 0.06488 copies of DNA per 2 µl reaction. Each concentration of synthetic target DNA was replicated eight times on a single qPCR plate, alongside three no-template controls (2 µl nuclease-free water in place of template DNA) and three positive controls (2 µl *N. rufogriseus* genomic DNA). The same qPCR reaction volumes were used as outlined above, and thermocycling conditions were as follows: 95°C for 3 minutes, 95 °C for 15 seconds, annealing at 58 °C for 30 seconds and 50 cycles, followed by 72 °C for 20 seconds. The curve fitting methods of Klymus et al., (2020) were applied using the provided R code and selecting the option “Best” to choose the best fitting model for calculating the LOD, and 35% CV threshold for calculating the LOQ.

### 2.2 Proof of Concept Experiments

Airborne eDNA was sampled in an open-air, controlled setting from *N. rufogriseus* wallabies at a captive park “EnkleDooVery Korna” housing males and females of different ages ranging from young-at-foot (juveniles permanently out of the pouch > 9 months) to adults. Wallabies were separated into groups of 1 – 5 individuals in various-sized paddocks spanning a total area of 5,705 m^2^ in Waimate, Canterbury, South Island, New Zealand (44°43’51.7” S 171°04’15.4” E). Two filter materials were tested for their effectiveness in capturing airborne eDNA. The first was a fibreglass air pocket bag filter (RS Components, New Zealand), classified as F7 grade under EN 779:2012, designed to capture fine dust particles ranging from 1–10 µm. The second was a fibreglass media roll typically used in spray painting booths (Filters Direct, New Zealand), rated as G4 grade under EN 779:2012, suitable for capturing coarse dust particles larger than 10 µm. Filter media were prepared for fieldwork in a dedicated eDNA pre-PCR laboratory. The air pocket bag was removed from the steel frame, the woven backing was peeled off and discarded, and the remaining filter media (∼3 mm thick) was cut to size using sterilised scissors. The fibreglass media roll was pulled into single layers (each layer ∼10 mm thick) and cut to size using sterilised scissors. Filter materials were each cut into 16 cm by 16 cm squares. Each side of the material was exposed to UV light for 30 minutes, then filters were stored in individual, sterile Ziploc bags.

Two air collection methods were tested for capturing airborne environmental DNA, an active and a passive collection approach. Active samplers were composed of a 12 V axial fan (IP55 model; Jaycar Electronics) mounted inside a custom 3D-printed plastic case, with the fan intake positioned approximately 3.5 cm from the filter material (Figure S1). The fan was powered by a battery pack containing four 1.5 V AA batteries. A single piece of either G4 or F7-rated filter material was secured over the front of the plastic frame using a sterile rubber band. The passive method used the same plastic case and filter setup, but without the fan or battery pack, relying solely on ambient air movement for DNA capture. Airflow rates from the battery-powered fan setup were measured as follows using a handheld anemometer (Uni T, UT363BT): 72.98 m³/h without a filter, 66.49 m³/h with a G4-rated filter attached, and 56.20 m³/h with an F7-rated filter attached (Table S2).

Air sampling was conducted on the 3^rd^ – 4^th^ August 2024 during the Austral Winter at the captive wallaby park with 23 *N. rufogriseus* wallabies in total. Active airborne eDNA samplers were housed in 6.4 L plastic containers (190 x 193 x 285 mm; Ezy Storage) mounted 1.5 metres above ground level using pull tie-down straps secured directly to the enclosure fencing (Figures 1A, 1B). A rectangular cutout (16 cm x 6 cm) on the front face of each container allowed unobstructed airflow directly onto the filters. Passive airborne eDNA samplers were attached directly to the enclosure fencing at a height of 1.5 metres using two sterile cable ties per unit (Figures 1A, 1B). Sampling took place within a single enclosure (26.5 metres x 14.5 metres) containing five adult wallabies, at five designated locations within the enclosure (Figure 2A).

**Figure 1.**
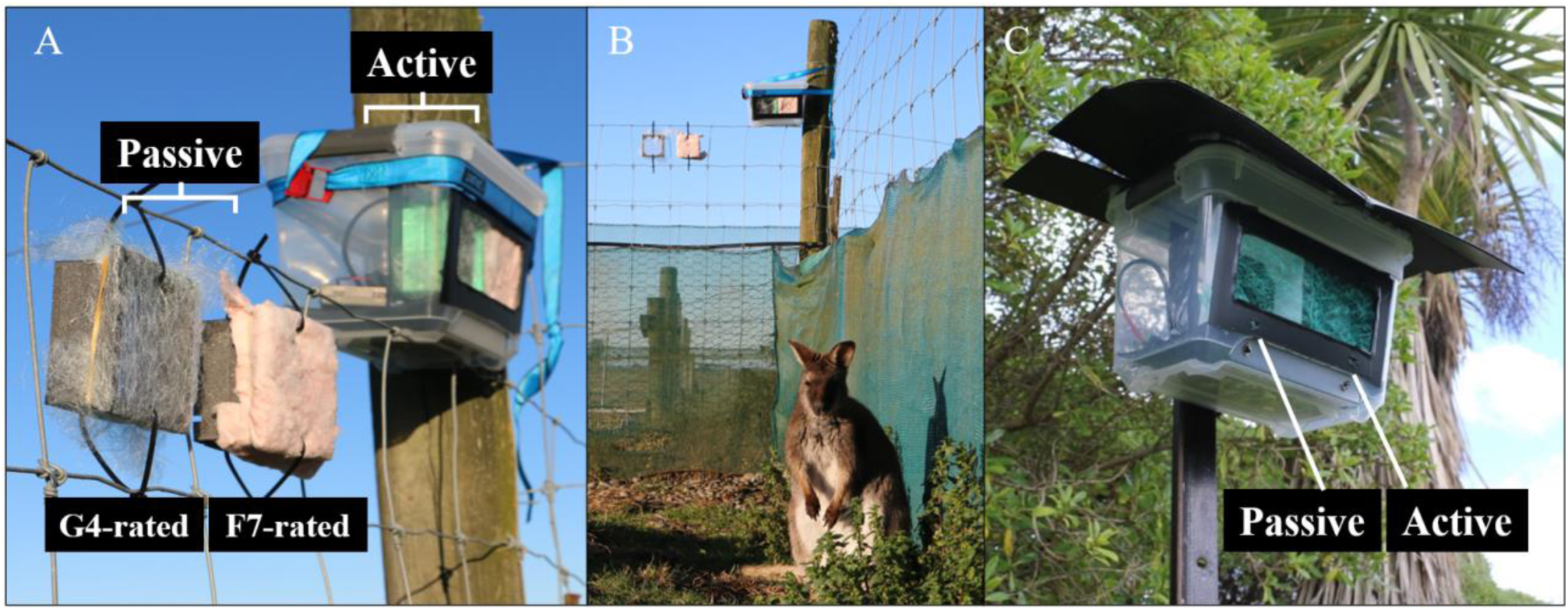
Airborne environmental DNA (eDNA) sampling approaches at Enkledoovery Korna Captive Wallaby Park, Waimate, South Island, New Zealand. Sampling employed two types of filter materials (G4-rated and F7-rated) and two collection methods (active and passive). **A, B)** Airborne sampler designs used in proof-of-concept experiments. **C)** Free-standing sampler used in distance experiments, featuring G4-rated material mounted on a metal stake with a corflute roof structure.

**Figure 2.**
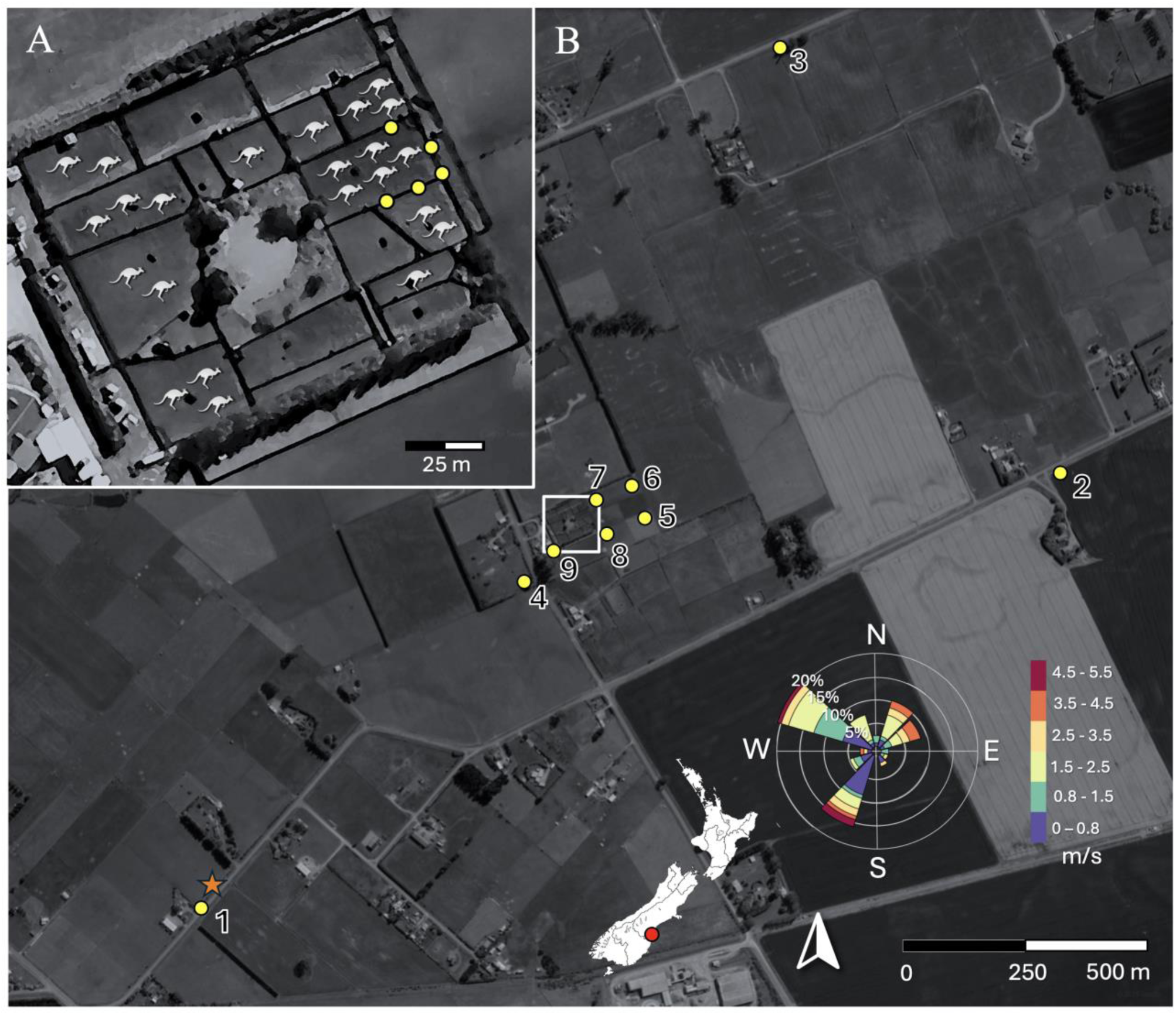
Airborne eDNA sampling sites (yellow) in relation to the Enkledoovery Korna Captive Wallaby Park, Waimate, New Zealand, during proof-of-concept experiments 3^rd^ – 4^th^ August 2024, and distance experiments 7^th^ – 12^th^ October 2024. **A)** The number and arrangement of wallaby individuals pictured during the proof-of-concept experiments. **B)** Average wind conditions, including frequency of wind direction and wind speed measured in metres per second (m/s), for the five sampling periods of the distance experiments. The orange star depicts the location of the weather station for the duration of the distance experiments.

At each location, two active and two passive air samplers were deployed, with each pair of active and passive samplers including both a G4-rated and an F7-rated filter. A total of 20 samples were deployed, with 10 for each filter type and each collection method. Each component of the air samplers (plastic containers, 3D-printed plastic cases, fans) were cleaned with 10% bleach (v/v) and stored in sterile Ziploc bags prior to sampling, then assembled on-site on a fold-out table disinfected with 10% bleach, while wearing sterile gloves. Air samplers were exposed for 23 – 24 hours (maximum sampling time of 24 hours 7 minutes, minimum 23 hours 15 minutes), covering the dusk and dawn periods where *N. rufogriseus* are known to be most active (Hickling & Day, 2024). Following sampling, filter housings were disassembled individually while wearing sterile gloves. Filters were removed using sterile tweezers, folded in half with the exposed side facing inward, and individually stored in sterile Ziploc bags. Between each disassembly, the fold-out table and tweezers were disinfected with 10% bleach, and new sterile gloves were worn. Samples were kept in a cooling box with ice for up to six hours before being transferred to –20 °C for long-term storage until extraction.

### 2.3 Distance Experiments

Airborne sampling was conducted on the 7^th^ – 12^th^ October 2024 during the Austral Spring at varying distances from the captive wallaby park with 27 *N. rufogriseus* wallabies in total. Fibreglass media roll (Filters Direct, New Zealand), rated G4 under EN 779:2012 (particle capture >10 µm), was selected for its demonstrated effectiveness in previous proof-of-concept experiments. Both active and passive collection methods were utilised to capture airborne eDNA. The same materials and field sampling approach were employed (see Section 2.2). However, the plastic containers housing the air samplers were mounted on metal stakes to position them approximately 1.5 metres above ground, allowing deployment in various open-air settings. Additional corflute “roof” structures were included on the lids of the plastic containers to protect the filters from exposure to rainfall (Figure 1C). Air sampling was conducted at nine locations; specifically, three sites each at distances of 10 metres, 100 metres, and 1000 metres from the captive wallaby park (Figure 2B). The selection of air sampling sites and the orientation of samplers were based on prevailing wind directions in the Waimate District, which are predominantly from the southeast and northwest, to maximise air flow.

At each location, one active and one passive air sampler were deployed, side-by-side inside a 6.4 L plastic container (Figure 1C). Air samplers were exposed to open air for durations ranging from a minimum of 23 hours to a maximum of 26 hours and 31 minutes. The experiment was repeated independently five times over five consecutive days, resulting in a total of 45 active and 45 passive field replicates, with 15 replicates of each collection method at each distance. To minimise the risk of DNA contamination between locations and sampling periods, sites were visited in the same order, beginning at the most distant sites (1 kilometre) to the closest sites (10 metres) in relation to the wallaby park. After each sampling period, sampler components were disassembled on a sterilised fold-out table while wearing sterile gloves and disinfected by soaking in 10% bleach for 10 minutes, followed by a rinse with distilled water. Components were then reassembled for the next use. Field personnel wore clean clothing each day, sterile gloves were changed and the fold-out table was disinfected between handling uncleaned and cleaned equipment and between every site. Local meteorological conditions, including temperature, humidity, wind speed, wind direction, and rainfall were obtained at hourly intervals using a digital weather station (IC-XC0432, Digitech) facing due south, held 1.5 metres above ground, at a site 1 kilometre from the captive wallaby park (Figure 2B).

### 2.4 DNA Extraction

DNA extraction was carried out in a dedicated eDNA pre-PCR laboratory, with several measures to reduce risk of contamination, including a unidirectional workflow, the wearing of hairnet, coveralls, two layers of medical gloves, allocated footwear, decontamination of surfaces and equipment using 10% bleach, and exclusive use of filter tips. DNA was extracted using the DNeasy Blood & Tissue Kit (QIAGEN, USA) following the manufacturer’s instructions, with some modifications outlined in detail below. Using sterile scissors, the F7 and G4-rated fibreglass filters were cut into smaller pieces, taken from the centre (G4-rated: 6 cm x 6 cm; F7-rated: 4 cm x 4 cm), then each piece was halved (G4-rated: 3 cm x 3 cm; F7-rated: 2 cm x 2 cm) and placed into separate 2 mL Eppendorf LoBind tubes using sterile forceps. After digestion, the corresponding halves of each air filter sample were recombined onto a single spin column. Filters were digested overnight at 56 °C with constant agitation at 1000 rpm. Each digestion tube contained 720 µl ATL buffer and 80 µl Proteinase K. Modified volumes of AL buffer and ethanol (600 µl each per sample) were used during extraction. DNA was eluted twice with 40 µl AE buffer per elution, each incubated at 37 °C for 15 minutes, resulting in a total elution volume of 80 µl. In addition to extracting field samples, negative filter controls were included, composed of sterilised air filter materials exposed to UV light for 30 minutes on both sides (n = 2 per filter type per experiment). Additionally, one negative extraction control was included for every 23 field samples to detect any potential contamination in the extraction reagents.

### 2.5 qPCR Analysis

All qPCR reactions were set up in a dedicated eDNA pre-PCR laboratory separate from the eDNA extraction laboratory, following the same protocols to reduce contamination risk as outlined above (Section 2.4). Each field sample was run through qPCR in triplicate, using the same reaction volumes and conditions as outlined above (Section 2.1.1). Field-collected samples were analysed alongside multiple negative controls, including negative extraction controls, negative filter controls, and PCR negative controls (n = 3 per plate). A standard curve was also included on each plate, consisting of a 10-fold dilution series of known target DNA copy numbers ranging from 64,880 -0.06488 copies per 2 µl reaction, with each standard concentration run in triplicate, using synthetic, double-stranded oligonucleotides of the *N. rufogriseus* MT-ND2 gene, consisting of the 106 bp PCR product flanked by additional 20 bp sequences on either end (total length 146 bp) inserted into a research-grade DNA plasmid (pUC57 vector). A field sample was considered positive for *N. rufogriseus* wallaby DNA if it met the following criteria: 1) the amplification curve crossed the threshold for detection within 40 cycles, 2) uniform curve morphology, 3) no detection observed in negative control samples. Samples that were detected below the limit of detection (LOD) were considered true positives if these same conditions were met (Hunter et al., 2017; Klymus et al., 2020). A single positive detection out of three technical replicates was considered a positive result, with detection in more than one replicate improving confidence. To test for the presence of inhibitors, a subset of 10 field samples were run through qPCR in neat and diluted concentrations with UltraPure water (1:10). Samples were considered inhibited if amplification of the neat concentration showed a delay compared to a 1:10 dilution. Positive detections were confirmed by Sanger sequencing to verify species identity. Post-PCR products were purified using a PALL filter plate protocol and prepared for bi-directional Sanger sequencing in a dedicated post-PCR area. Sanger sequencing was carried out by Genetic Analysis Services, Department of Anatomy, University of Otago (Dunedin, New Zealand).

### 2.6 Statistical Analyses

All statistical analyses were conducted in R version 4.5.1 (R Core Team, 2025). Significance was evaluated at the *p* ≤ 0.05 level. A field sample was considered positive if any of its three technical replicates yielded a positive amplification result. Detection outcomes were recorded as either a positive amplification result (1) or a negative result (0). Kruskal-Wallis rank-sum tests were conducted to assess differences in cycle quantification (Cq) values, and DNA quantity (DNA copies/µl) across sample collection methods (active versus passive collection), filter types (F7 versus G4-rated filters), and distance from the source. Where significant differences were found, post-hoc pairwise comparisons were performed using Dunn’s test with Bonferroni correction. To address issues of separation in the data, particularly the strong association between filter type and positive detections, Firth’s penalised logistic regression model was fitted to the data using the *logistf* package v1.26.1 (Heinze et al., 2025). Detection rates were also compared between pairs of actively and passively collected air samples from the same sites and on the same days using Wilcoxon rank-sum tests. For the distance experiments, McNemar’s test was also applied to assess whether the probability of detection being positive for active samplers but negative for passive samplers differed significantly from the reverse scenario.

Detection probabilities were calculated for each distance category (10, 100, and 1000 metres) and sample collection method (active and passive) as the proportion of positive samples. These probabilities were then used to estimate the number of field replicates required to achieve a 95% probability of at least one positive detection under two scenarios: (1) a source population of 27 wallabies, which was the number of wallabies present within the boundaries of the entire captive wallaby park during the distance experiments, and (2) a single wallaby. Generalised linear models (GLMs) were applied with a binomial distribution to model detection outcomes as binary variables (positive = 1, negative = 0). These models allowed an assessment of the fixed effects of distance, collection method, and environmental variables on detection probability. Fixed environmental variables were; **Temperature**: The average air temperature during the sampling period, measured in degrees Celsius. **Humidity**: The average relative humidity percentage during the sampling period. **Wind Speed**: The average wind speed during the sampling period, measured in metres per second. **Wind Direction**: an average of the direction from which the wind was blowing from, expressed in degrees from true north (0° indicating wind from the north) for the sampling period. Compass directions were converted to degrees, from degrees to radians, and transformed from a linear variable to *sin* and *cos* components. **Rainfall**: The total amount of precipitation that occurred during the sampling period, measured in millimetres. **Site Direction**: The compass direction that the sampler was facing at each site, expressed in degrees from true north, with data transformed the same as for wind direction. Active and passive airborne sampler data were combined in a single model, with the collection method included as a fixed effect to evaluate its impact on detectability. Separate models were also run for active and passive datasets to identify collection-specific effects. To choose the best model, backward model selection was performed with the *dredge* function from the MuMIn package v1.46.0 (Bartoń, 2022), starting from the most comprehensive model including all predictors. Models were ranked by Akaike Information Criterion corrected for small sample size (AICc), and the best-fitting models were chosen based on the lowest AICc values. Wald tests (Pr(>|z|)) were used to determine the significance of fixed effects, including distance, sampler type, and environmental variables. Models were considered significant if *p* < 0.05 and 95% bootstrap confidence intervals excluded zero.

A correlation matrix was generated to examine relationships among predictor variables, particularly environmental factors, and to assess the significance of these correlations. To address potential multicollinearity among environmental predictors, Principal Component Analysis (PCA) was performed on environmental variables including wind speed, wind direction, temperature, humidity, and site direction. The resulting principal components (PCs) were then used as predictors in a logistic regression model to examine their collective influence on detection probability.

A Bayesian logistic regression model in the *brms* package v2.22.0 (Bürkner, 2017) was also applied to include the effects of sampling site and sampling day to account for spatial and temporal variation. The response variable was detection outcome (1 = detected, 0 = not detected), modelled using a Bernoulli distribution with a logit link function. Predictor variables included those from the GLM. The model was run with four Markov Chain Monte Carlo (MCMC) chains, each with 4,000 iterations (1,000 warm-up), yielding 12,000 post-warmup samples. Convergence was assessed using R-hat statistics and effective sample sizes.

## 3 RESULTS

### 3.1 Primer Design and Validation

For the targeted *N. rufogriseus* assay (Table 1), co-amplification of nontarget taxa was expected to be low, with multiple mismatches occurring in the forward, reverse, and probe regions. However, two macropods that do not overlap in range with the target species in New Zealand had either a single mismatch to the probe region (*Petrogale xanthopus*) or zero mismatches to the probe region (*Notamacropus agilis*) (Table S3), which could be tested further if the assay were applied beyond New Zealand. Primers successfully amplified the target DNA across the full range of annealing temperatures, with no amplification observed in no-template negative controls. Based on Cq values and the intensity of banding patterns on the gel, the optimal annealing temperature for the targeted assay was determined to be 58 °C. Laboratory validation using genomic DNA extracts confirmed that the DNA of the target wallaby species was successfully amplified by the target assay, and that the DNA of closely related/co-occurring mammalian species were not amplified. The assay had a LOD of 9 copies of DNA per 2 µl reaction, and a LOQ of 23 copies of DNA per 2 µl reaction. The PCR efficiency was calculated as 90.91% (Figure S2; Table S4).

**Table 1.**
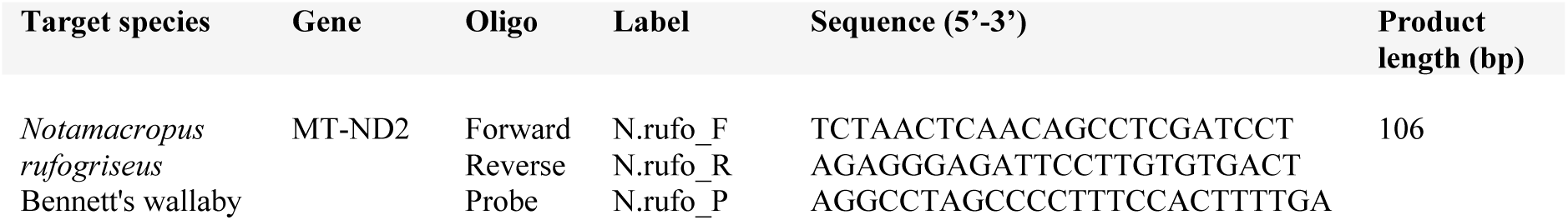
Primer and probe sequences designed to amplify a short region of the MT-ND2 gene in Bennett’s wallaby *Notamacropus rufogriseus*, showing the oligonucleotide sequences and PCR product length (base pairs).

### 3.3 Proof of Concept Experiments

Of 20 airborne eDNA samples collected in the captive wallaby enclosure, 14 (70%) tested positive. Positive detections were confirmed by Sanger sequencing. Sequences displayed ≥ 98% similarity with *N. rufogriseus* sequences in NCBI. Of 14 positive samples, 10 of 14 (71.4%) were collected using the G4-rated filter material. Furthermore, 9 of 14 samples were collected using the active collection method, whereas 5 were passively collected. All passive samples utilising the F7-grade collection material failed to amplify. After fitting Firth’s penalised logistic regression model, G4-rated filter material had a significant positive effect on detection probability (*p* = 0.00105), while passive samplers had a significant negative effect on detection probability (*p* = 0.015). However, pairwise Wilcoxon rank-sum tests found no significant difference in detection rates between actively and passively collected air samples from the same sites.

Quantitative comparisons using Kruskal-Wallis rank-sum tests showed that actively collected samples recovered an estimated 10.45 more DNA copies/µl on average compared to passively collected samples, although this was not a statistically significant difference (*p* = 0.4132). The Cq values did not differ significantly between collection methods (difference = 0.310 Cq units, *p* = 0.8348). In contrast, among actively collected samples, those using G4-rated filters had significantly lower Cq values than those with F7-rated filters (difference = -2.146 Cq units, *p* = 0.0138), and recovered 145.928 more DNA copies/µl, a significant difference in DNA quantity (*p* = 0.0138) (Figure 3).

**Figure 3.**
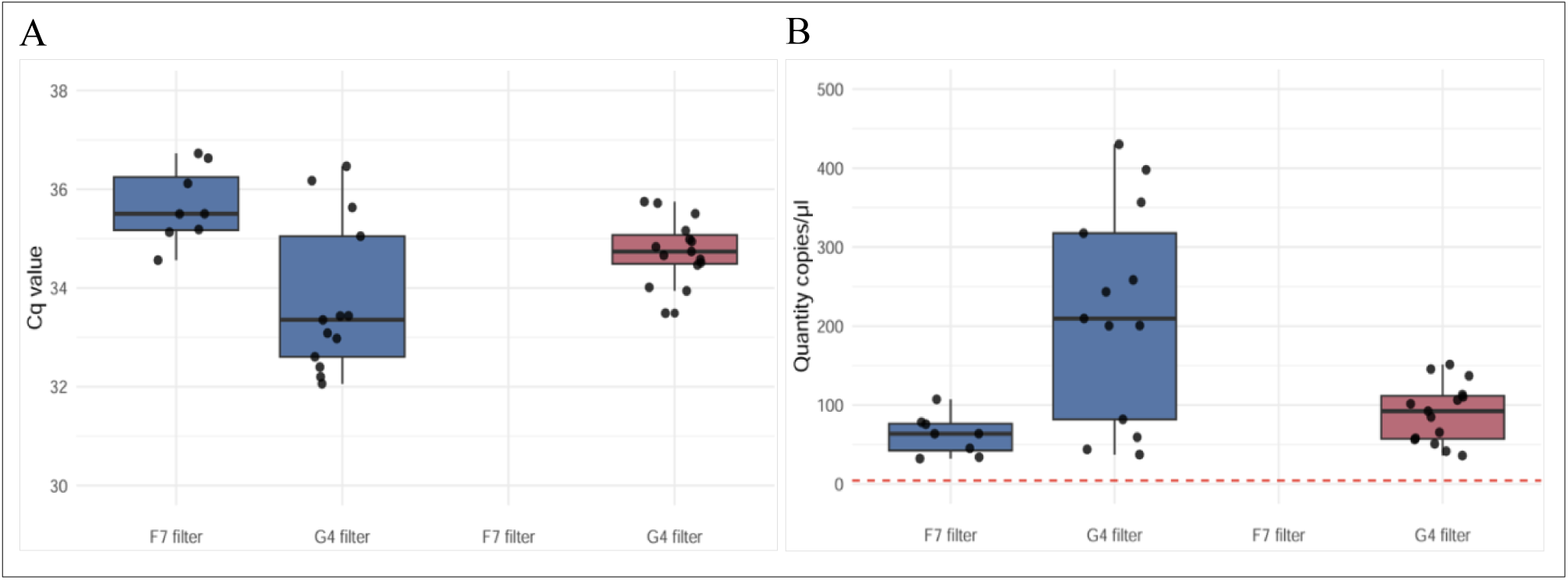
Airborne eDNA samples positive for *Notamacropus rufogriseus* DNA using two collection methods: active sampling (blue), and passive sampling (red), and two filter types: F7-rated and G4-rated airborne filters. **(A)** Cycle quantification (Cq) values calculated for each positive eDNA sample collected by airborne eDNA sampling, **(B)** DNA copy number (copies/µL) recovered from airborne eDNA sampling, with dashed red line indicating the limit of detection (LOD) for this assay; no samples fell below this threshold. Kruskal-Wallis rank-sum tests show a non-significant difference in DNA copies/µl or Cq value of samples by collection method (*p* = 0.4132; 0.8348), but a significant difference in DNA copies/µl and Cq value of samples collected using G4-rated or F7-rated filter types (*p* = 0.0138; 0.0138).

Samples were considered inhibited if amplification of the neat (undiluted) concentration of a sample showed a delay in amplification compared to a 1:10 dilution. Based on these criteria, no evidence of inhibition was detected. As such, samples were processed without an additional step for inhibition removal.

### 3.4 Distance Experiments

Of 88 airborne eDNA samples collected at various distances from the captive wallaby park, 27 samples (30.7%) tested positive for wallaby eDNA. Detection probability varied with distance. At 10 metres, 16 of 30 samples (53.3%) were positive; at 100 metres, 7 of 30 samples (23.3%) were positive; and at 1 kilometre, 4 of 28 samples (14.3%) were positive. Detection rates were higher in samples collected using the active collection method. Of 27 positive detections, 20 (74.1%) were actively collected. At 10 metres, the detection rate for active samplers was 64.4%, more than five times higher than the 11.1% positive detection rate observed for passive samplers. Similarly, at 100 metres, detection rates for active and passive samplers were 13.3% and 4.44%, respectively. However, at 1 kilometre, detection rates were low and comparable between collection methods (4.44% for active samplers; 6.67% for passive samplers) (Figure 4). With 27 captive wallabies acting as the source of wallaby DNA in this experiment, the number of airborne field replicates required to achieve 95% confidence of at least one positive detection was considerably lower than the number of replicates predicted for detecting a single wallaby (Table S5). Specifically, using active airborne samplers, only 3 field replicates would be needed at 10 metres, 21 replicates at 100 metres, and 66 replicates at 1000 metres. In contrast, for a single wallaby, the required number of active airborne field replicates increases substantially, with 79 replicates needed at 10 metres, 567 at 100 metres, and 1781 at 1000 metres.

**Figure 4.**
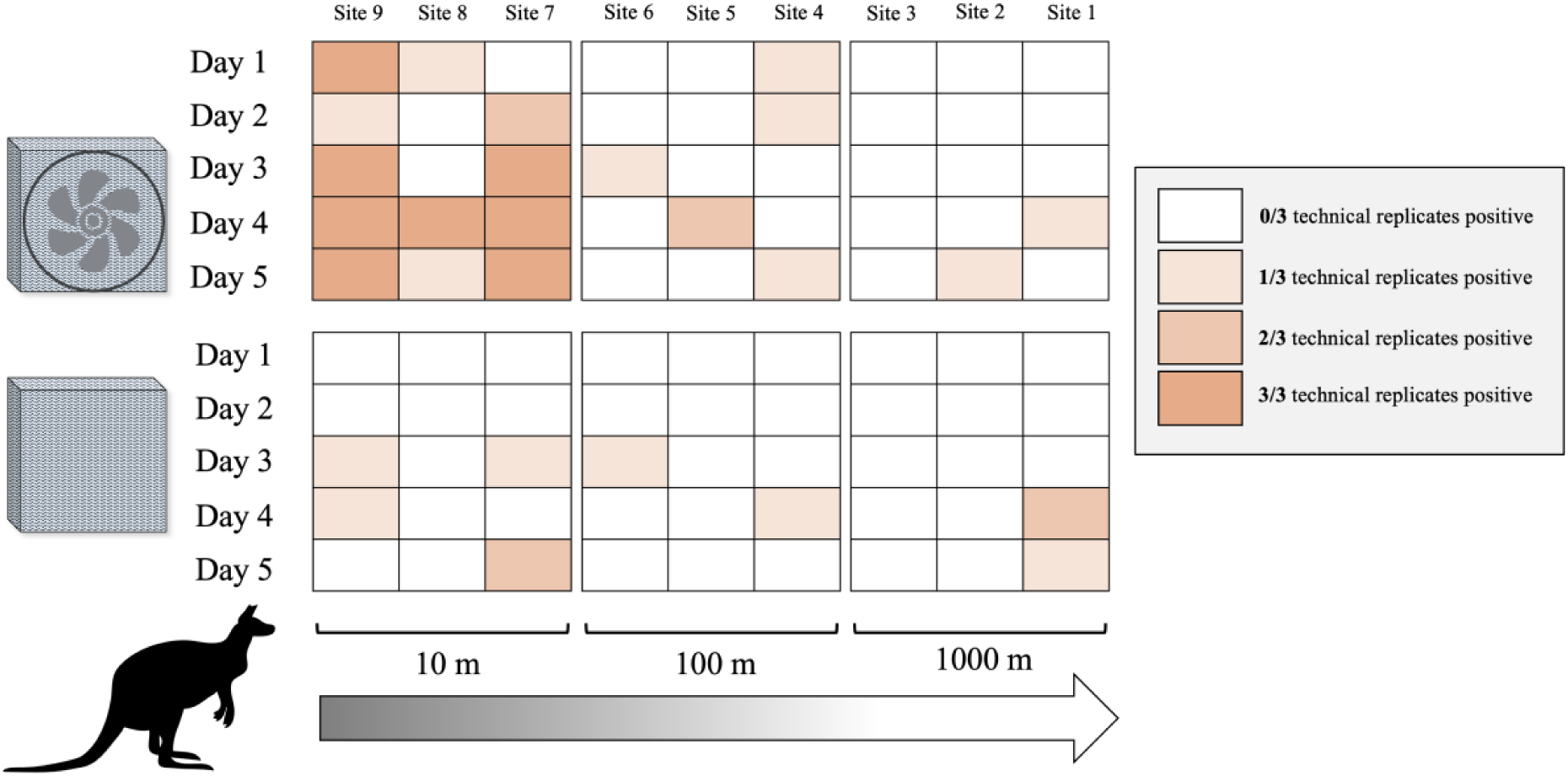
Frequency of positive airborne eDNA detections of *Notamacropus rufogriseus* obtained using two collection methods, active sampling (fan-powered) and passive sampling (no fan) at 10, 100, and 1000 metres from the source of wallaby DNA at the Enkledoovery Korna Captive Wallaby Park, Waimate, New Zealand, 7^th^ – 12^th^ October 2024.

Generalised linear models (Table S6) revealed that distance from the source had a strong and significant negative effect on detection probability in both the combined active/passive dataset (effect size = -1.0715, *p* = 0.00226), and the active dataset (effect size = -1.343, *p* = 0.00473), suggesting detection probability declines consistently with increasing distance from the source of wallaby DNA when using the active collection method. In contrast, distance was not retained in the best-fit model for passive collection, suggesting a weaker or inconsistent distance effect with this method. Bayesian logistic regression (Table S7) also found that for active samplers, distance negatively affected detection (estimate = -0.91, 95% CI: -1.73 to -0.10), whereas in passive samplers, distance effects were uncertain (estimate = -0.42, 95% CI: -1.53 to 0.61). Moreover, the GLMs demonstrated that passive samplers were significantly less effective than active samplers in detecting airborne wallaby eDNA (effect size = -1.6191, *p* = 0.00618). Similarly, using Bayesian logistic regression for the combined active/passive dataset, passive samplers had significantly lower detection odds (estimate = -1.30, 95% CI: -2.31 to -0.34), reflecting about 73% reduced detection probability. Additionally, Wilcoxon rank-sum tests comparing paired active and passive samplers deployed concurrently at the same sites and on the same days identified significantly higher detection rates for active samplers at several 10-metre sites, specifically, Site 9 (Days 1 and 5), Site 8 (Day 4), and Site 7 (Day 4) (*p* < 0.05). However, for sites located at distances beyond 10 metres, no significant differences in detection rate between collection methods were observed.

Environmental variables were measured consistently over the 5 sampling periods (Figs. S3-S6) and were evaluated for their influence on airborne eDNA detection probability. In the GLMs, average wind speed, wind direction, and humidity did not significantly affect detection probability and were excluded from all best-fitting models. In contrast, the variable *site_direction_sin*, which reflects airborne sampler orientation relative to the direction the wind is travelling from, was a significant predictor of detection probability in the combined active/passive dataset (*p* = 0.0023), as well as the active (*p* = 0.0397) and passive (*p* = 0.0275) datasets individually. Despite these significant effects, wide confidence intervals including zero indicate uncertainty regarding the strength and direction of this effect. Moreover, temperature had a significant negative effect on detection probability in the combined active/passive dataset (*p* = 0.0403, effect size = -0.578), suggesting that higher temperatures may reduce eDNA detectability. However, this result should also be interpreted cautiously due to wide confidence intervals. Bayesian logistic regression found that none of the environmental variables showed clear effects. However, random intercepts for site and day accounted for substantial variation in detection probability (Table S8), with odds of detection differing approximately 2-to 2.5-fold depending on location or sampling day, indicating important spatial and temporal heterogeneity not captured by the GLM.

Statistically significant correlations were observed between certain environmental variables, such as wind speed and temperature (r = -0.45, *p* = 8.88e-15) and wind speed and humidity (r = -0.47, *p* = 2.22e-16). To address multicollinearity, a PCA performed on the environmental variables from the combined dataset of active and passive samples identified three main components that together explained approximately 85% of the total variance in detection rates. Specifically, PC1 accounted for 43% of the variance and reflected meteorological conditions such as wind speed and wind direction; PC2 explained 25% of the variance and was strongly associated with site orientation; PC3 captured 17% of the variance and related mainly to temperature and humidity patterns. These three PCs were then used as predictors in a logistic regression model to evaluate their collective influence on airborne eDNA detection probability. The results indicated a statistically significant negative effect of PC2 (estimate = -0.42, p = 0.0149), whereas PC1 and PC3 showed no significant effect on detection probability. Both *sin* and *cos* components of site direction had strong negative loadings, suggesting that site directions pointing in a certain orientation were associated with a lower probability of detection.

Kruskal-Wallis rank-sum tests were conducted using Cq values and DNA copy number (copies/µl) as response variables to examine how DNA concentration varied with airborne collection method and distance from the source. For actively collected samples, distance was a significant predictor of both Cq values (*p* = 0.0075) and DNA quantity (*p* = 0.0374). In contrast, for passively collected samples, distance was not a significant predictor of either Cq values (*p* = 0.5159) or DNA quantity (*p* = 0.6244), indicating no clear trend in DNA yield with increasing distance (Figures S7, S8). Overall, positive samples that were passively collected exhibited significantly higher Cq values than active samples (*p* = 0.04). Passively collected samples also yielded lower DNA quantities (median ∼15 copies/µl less) compared to actively collected samples, though this was not significant (*p* = 0.073).

## 4 DISCUSSION

We successfully developed and validated a novel, species-specific probe-based qPCR assay for *N. rufogriseus*. This assay demonstrated high sensitivity, reliably detecting DNA at concentrations as low as 4.5 copies/µl, and in laboratory trials showed no cross-reactivity with closely related or co-occurring taxa. In a controlled field setting, airborne eDNA samples yielded positive results for *N. rufogriseus*. Several factors influenced airborne detectability, including filter material, distance from the source, collection method, and environmental conditions.

### 4.1 Effect of Filter Type

At close range to the source of wallaby DNA, coarse filters rated for capturing particles >10 µm were optimal, generating higher detection rates, greater DNA concentrations (DNA copies/µl) and lower Cq values, compared to filters that targeted finer particles (1–10 µm). These findings align with those of Bodawatta et al., (2025), who reported that filters capable of retaining a broader range of airborne particle sizes, including larger particles, were more effective for characterising terrestrial vertebrate communities. Moreover, a larger filter surface area enhanced their vertebrate DNA detectability. The substantially large filters used in this study (16 × 16 cm) may have improved capture of rare or sporadically distributed airborne eDNA. Observed differences in DNA yield between filter types may reflect variation in both eDNA capture efficiency and extraction efficiency. The coarse, G4-rated filters were thicker and more fibrous, potentially enhancing their ability to trap a broader range of airborne particle sizes. Their improved performance could be attributed to greater resistance to clogging, allowing for sustained airflow over extended sampling periods, or alternatively, more efficient capture of the airborne particle size range shed from terrestrial vertebrates such as wallabies at close range. Additionally, differences in DNA recovery may have contributed to the observed variation. The coarse filters appeared to release reagents such as lysis buffer more readily during extraction, whilst the finer filter material absorbed a larger volume of reagents during the overnight lysis step and retained a significant volume, potentially limiting DNA yield. Thus, the superior performance of coarse filters may result from a combination of factors, which could be disentangled by future studies incorporating internal positive controls to directly quantify DNA extraction efficiency.

### 4.2 Effect of Distance

Positive airborne eDNA detections were more likely when airborne eDNA samplers were positioned closer to the source of wallaby DNA. For active (fan-powered) sampling, distance emerged as the strongest predictor of detection probability, with a clear decline in both detection rates and DNA concentrations beyond 10 metres from the wallaby park. This is consistent with dilution and dispersal effects whereby increasing distance reduces eDNA concentration beyond detectable limits (Jerde et al., 2016; Shogren et al., 2017), likely reflecting a combination of rapid dilution, fast degradation of airborne particles, and limited transport of larger DNA-bound particles, which may settle quickly and travel only short distances, constraining detection range. Consequently, detection at greater distances was sporadic, consistent with reports of stronger signals near source animals and reduced detection with increasing distance (Clare et al., 2022; Roger et al., 2022; Lynggaard et al., 2024; Jager et al. 2025). While detectability was highest when samplers were positioned closer to the source, long-distance dispersal remained possible up to 1 kilometre distances, for both active and passively collected samples. Similarly, Jager et al. (2025) reported positive airborne eDNA detection using a passive sampler 515 metres from the source within 24 hours of deployment. This dispersal potential underscores the value of airborne eDNA for landscape-scale surveillance, particularly in the context of managing highly mobile terrestrial species.

### 4.3 Effect of Airborne Collection Method

Both active and passive collection methods successfully detected airborne wallaby eDNA in a natural landscape, at distances of up to 1 kilometre from the closest known source. However, active samplers consistently outperformed passive samplers at close ranges, specifically, at distances of 0 to 100 metres. This is potentially due to their ability to process larger air volumes over short deployment periods. Bodawatta et al. (2025) similarly found that higher air volumes filtered by 12V fan-driven active samplers improved detection of terrestrial vertebrates compared to passive samplers. However, at distances greater than 100 metres, we observed no difference in detection rates between the two airborne collection methods, suggesting both may approach a shared efficiency threshold where detections become more dependent on sporadic eDNA dispersal via wind or other stochastic factors.

Passive and active sampler performance was highly variable, with limited replication in positive samples consistent with prior reports of patchy, low abundance of airborne vertebrate eDNA in open-air environments (Lynggaard et al. 2024). This variability underscores the importance of strong replication at both field and technical levels. In contrast to our findings, Jager et al. (2025) reported greater effectiveness of passive sampling when samplers were deployed for extended durations (96 hours), compared to brief (10-minute) active sampling intervals. Their higher detection rates for passive samplers were likely driven by greater cumulative air capture, suggesting that deployment times longer than the 24 hours used in this study may substantially enhance passive sampler performance.

### 4.4 Effect of Environmental Conditions

Environmental conditions may have also impacted airborne detectability in our experiments. Higher temperatures were associated with reduced airborne eDNA detection rates, which may reflect both environmental and biological mechanisms. Higher temperatures are known to accelerate DNA degradation (Jo et al., 2019; Naef et al., 2023). Temperature may also influence airborne detection rate through its effect on animal behaviour. Bennett’s wallabies are crepuscular, with peak activity at dawn and dusk (Hickling & Day, 2024), which are periods typically associated with cooler temperatures. Increased activity when it is cooler may result in greater DNA shedding, thereby increasing airborne eDNA availability. Moreover, sampler orientation appeared to be a significant predictor of detectability. Airborne samplers, particularly passive samplers, are likely heavily reliant on wind direction aligning with the direction of the filter in order to capture particulates (Waza et al., 2019; Johnson et al., 2023). These effects may also be modulated by local topography, which can funnel or disperse DNA-carrying air masses. Importantly, substantial variation in detection was explained by differences among sampling locations and days, indicating that even at fine scales, environmental heterogeneity influences airborne eDNA capture.

### 4.5 Caveats and Future Recommendations

Despite the patterns observed, several sources of uncertainty remain. While unlikely, the presence of other wallabies, or sources of wallaby DNA outside of the captive wallaby park, cannot be ruled out and may have contributed to detections at unexpected distances, potentially confounding the distance-based analysis. Unmeasured variables such as vegetation density, topography, and airborne sampler height above ground may also have influenced airborne eDNA dispersal, capture, or degradation. In addition, relatively small sample sizes may have reduced our statistical power, thus, greater sampling replication in future may increase our ability to clarify relationships between detectability and environmental variables. Whilst airborne eDNA demonstrated high sensitivity for detecting wallabies in captive settings, it remains untested whether these methods could sufficiently detect varying wallaby densities in natural environments where long-range dispersal is possible. Additionally, the degradation rate of airborne eDNA in open-air environments is currently unknown. This information will be critical for informing surveillance programmes, particularly in determining optimal deployment durations for samplers.

### 4.6 Implications for Management

To support the management of invasive species, eDNA assays must be both validated and practically deployable. Our targeted *N. rufogriseus* assay meets Level 5 (operational) on the five-level validation scale proposed by Thalinger et al., (2021). However, translating this assay for effective management schemes requires determining adequate field and technical replicate numbers to detect weak or sporadic signals, establishing clear thresholds for management responses, and identifying contexts where airborne eDNA is best suited for surveillance of the target (Morisette et al., 2021; De Brauwer et al., 2023). The number of field replicates we calculated as necessary to achieve 95% confidence in at least one positive detection highlights how detection efficiency decreases sharply with fewer animals and at greater distances from the target, requiring a much greater sampling effort to reliably detect DNA. Despite this, airborne eDNA has the potential to be cost-effective when integrated into a tiered surveillance framework. We propose that airborne eDNA serves as a broad-scale, low-cost screening tool to identify areas that warrant more detailed, labour-intensive follow-up investigations. Traditional surveillance techniques are likely to be most economical in areas where the likelihood of target presence is moderate to high, whereas airborne eDNA is likely to be well-suited for monitoring terrestrial areas with very low expected target densities, where conventional methods may be inefficient or prohibitively expensive.

Although logistical challenges currently limit the large-scale deployment of our airborne eDNA samplers, these issues could be mitigated through the development of more portable materials and simplified field protocols. Additionally, incorporating a mechanism that allows the sampler to rotate and capture air from all directions is likely to greatly enhance detection efficiency, given that we identified sampler orientation as a significant factor influencing detectability. A key outcome of this study is the significant reduction in material costs achieved by substituting previously utilised airborne filter materials such as F7-grade fibreglass pocket filter bags or Filtrete 1900 MERV13 with a fibreglass media roll. The media roll used in this study is priced at approximately $5.33 USD per metre, yielding enough material to supply 100 airborne sampling units, thereby enabling the large scalability that would be required for successful surveillance programmes.

In summary, we developed a sensitive, species-specific qPCR assay for detecting airborne eDNA of the invasive terrestrial vertebrate *N. rufogriseus*, demonstrating its potential as a tool for broad-scale surveillance. Our results show that using coarse filters and active collection methods significantly improves airborne detection probability of wallabies, as well as sampling at close range, though long-range detections remain possible. Environmental variables such as temperature and wind direction relative to sampler orientation likely influence airborne eDNA dynamics. While further work is needed to refine deployment strategies and assess assay performance in natural settings, especially at low target densities, these findings suggest that airborne eDNA offers a promising, cost-effective complement to existing terrestrial species monitoring tools. With continued optimisation, airborne eDNA could enhance early detection and support targeted management of low-density terrestrial species.

## Supporting Information-Targeted Airborne eDNA Detection of Pest Wallabies

**Figures S1-S8**

**Tables S1-S3, S5-S11**

**Figure S1.**
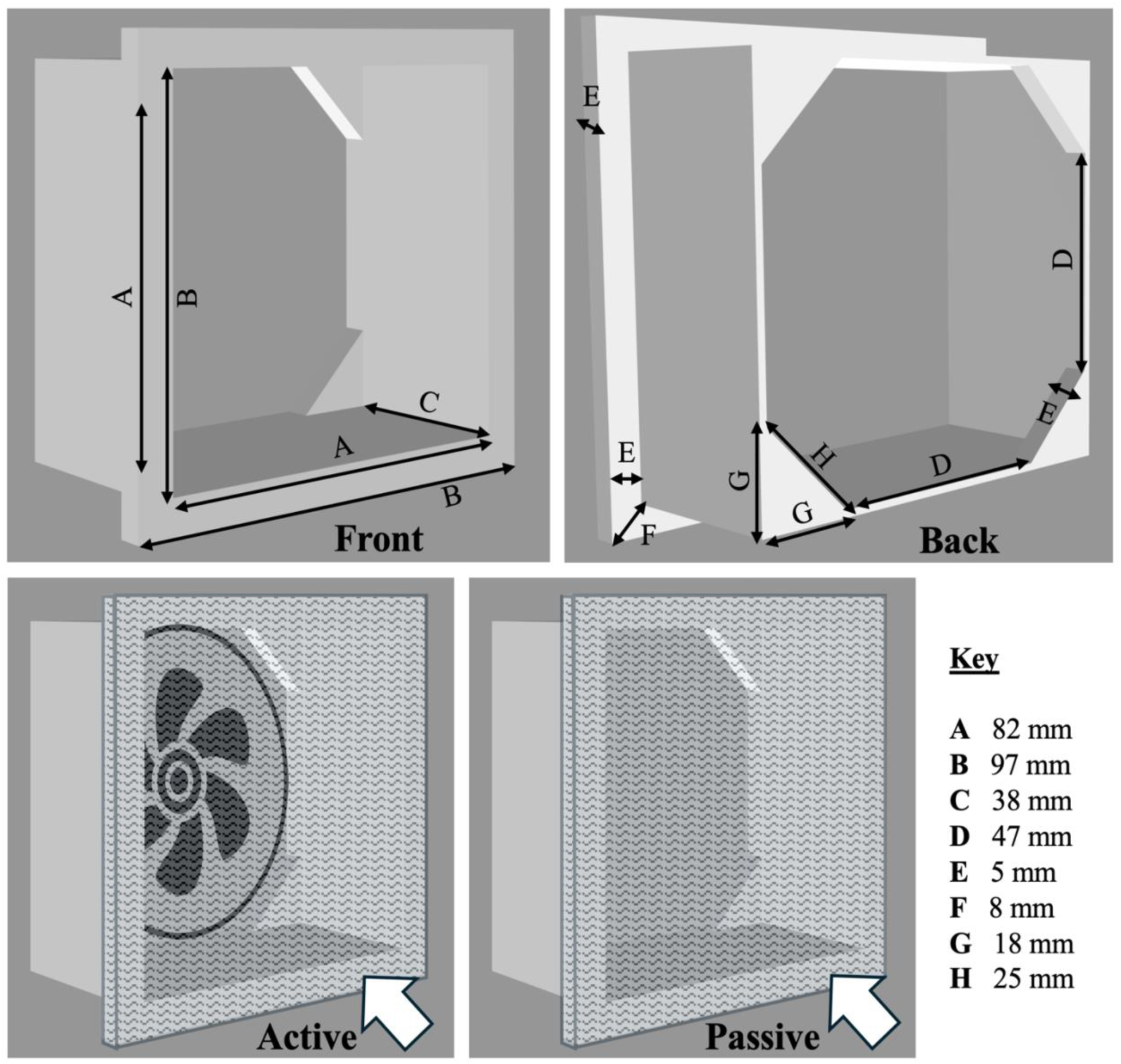
Airborne eDNA sampler design depicting the dimensions of the plastic 3D-printed frame, assembly of the active (fan-powered) and passive (no fan) collection system, and direction of wind (white arrows) in relation to the position of the sampler.

**Figure S2.**
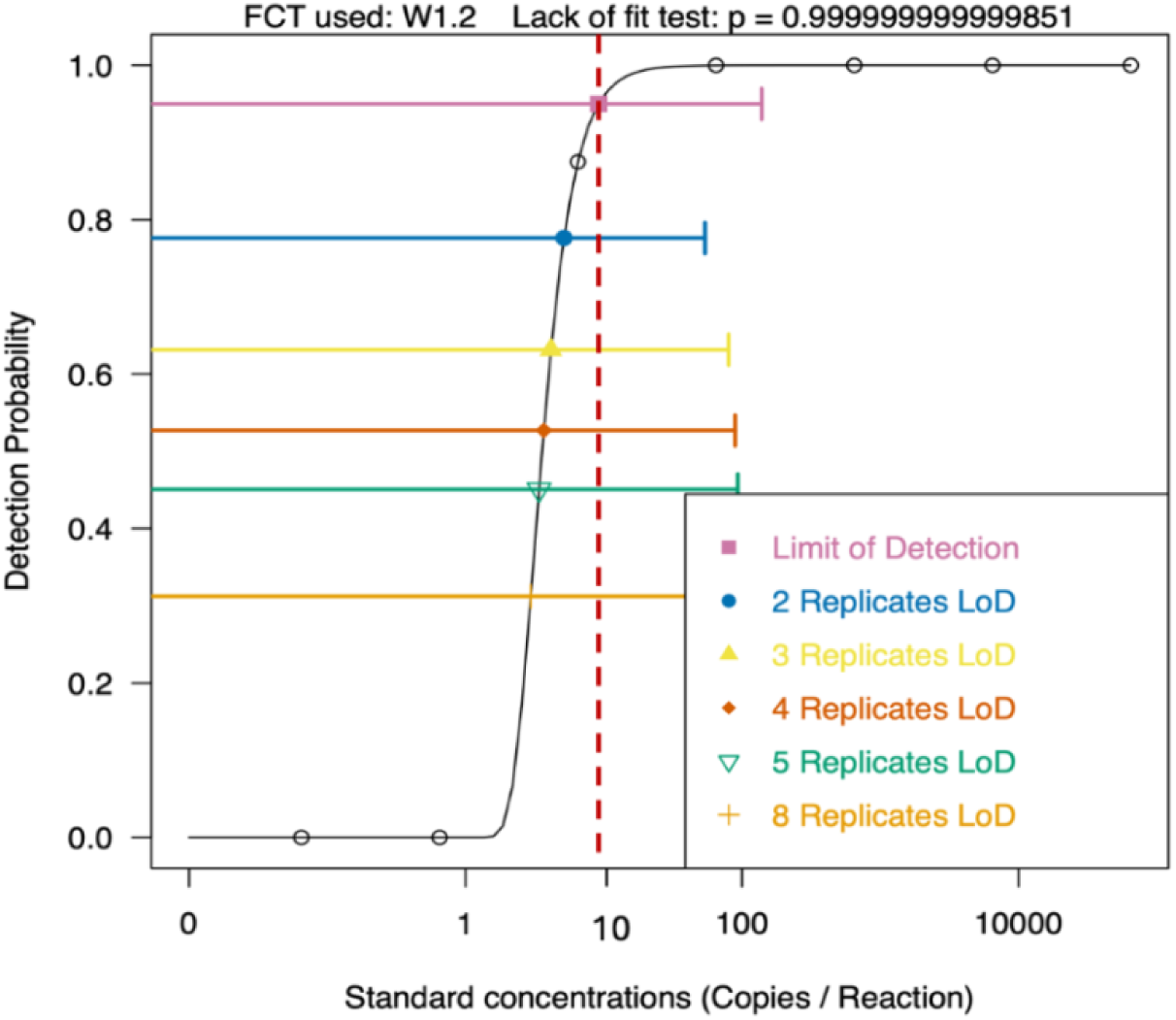
Limit of Detection (LOD) calculated for the Bennett’s wallaby assay measured using quantitative PCR (qPCR) at 2 µl reaction volume, using 146 bp region of the MT-ND2 gene in Bennett’s wallaby *Notamacropus rufogriseus* within a pUC57 plasmid (GenScript), total length 2856 bp.

**Figure S3.**
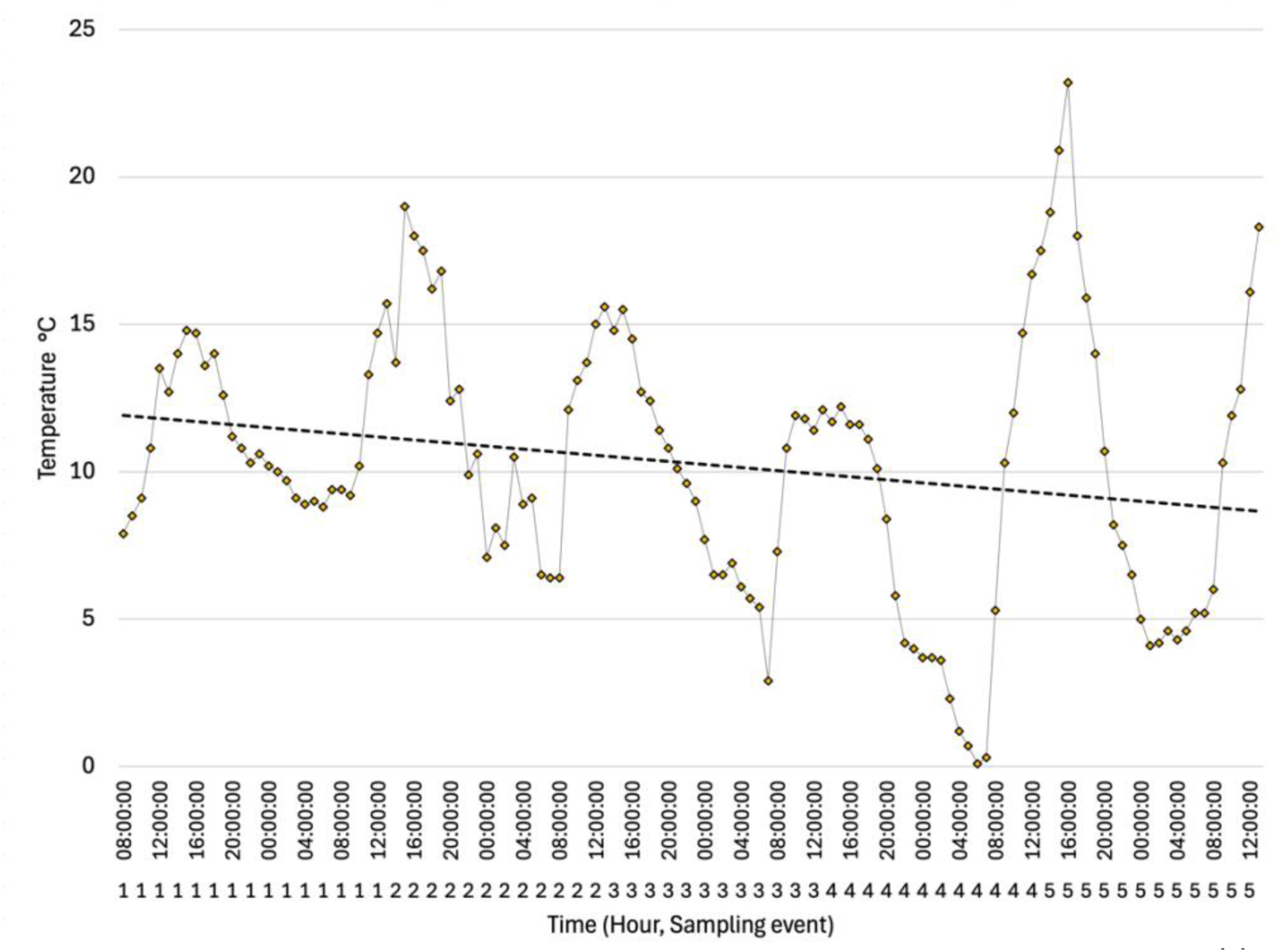
Temperature (°C) across five sampling periods of distance experiments 7^th^ – 12^th^ October 2024 Waimate, New Zealand, measured by local weather station set up 1 kilometre distance away from the captive wallaby park EnkleDooVery Korna.

**Figure S4.**
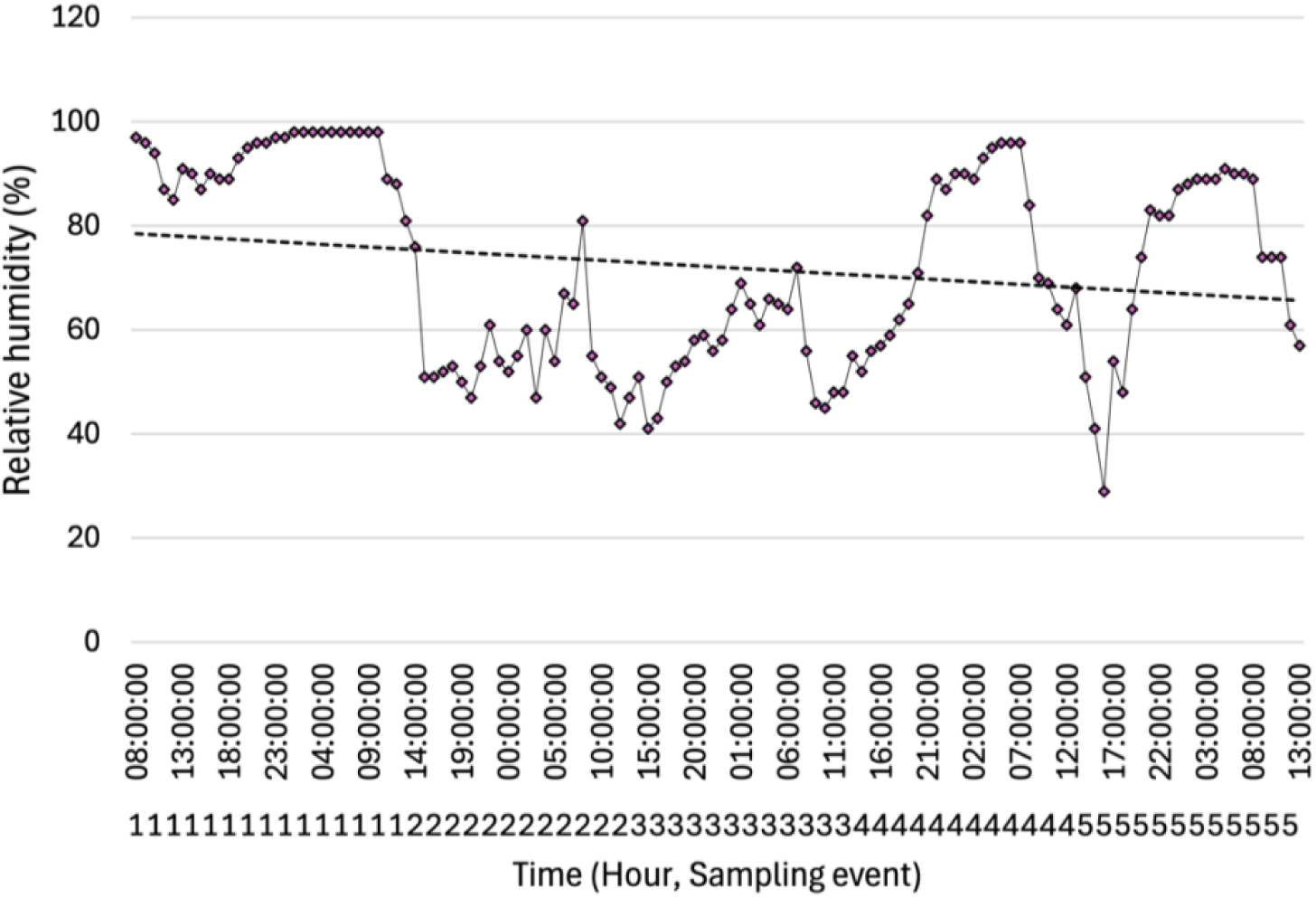
Relative humidity (%) across five sampling periods of distance experiments 7^th^ – 12^th^ October 2024 Waimate, New Zealand, measured by local weather station set up 1 kilometre distance away from the captive wallaby park EnkleDooVery Korna.

**Figure S5.**
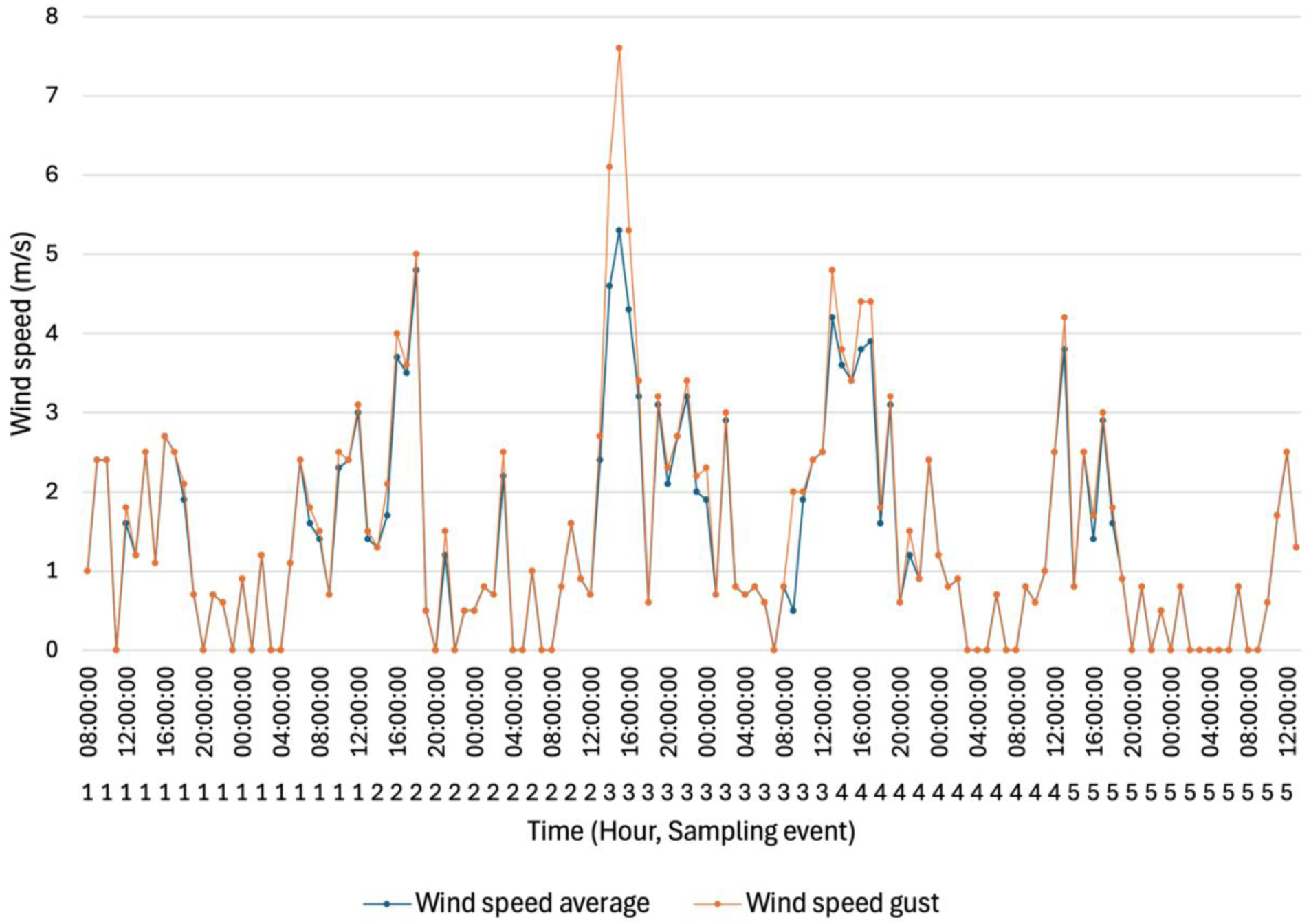
Wind speed average and wind speed gust (m/s) across five sampling periods of distance experiments 7^th^ – 12^th^ October 2024 Waimate, New Zealand, measured by local weather station set up 1 kilometre distance away from the captive wallaby park EnkleDooVery Korna.

**Figure S6.**
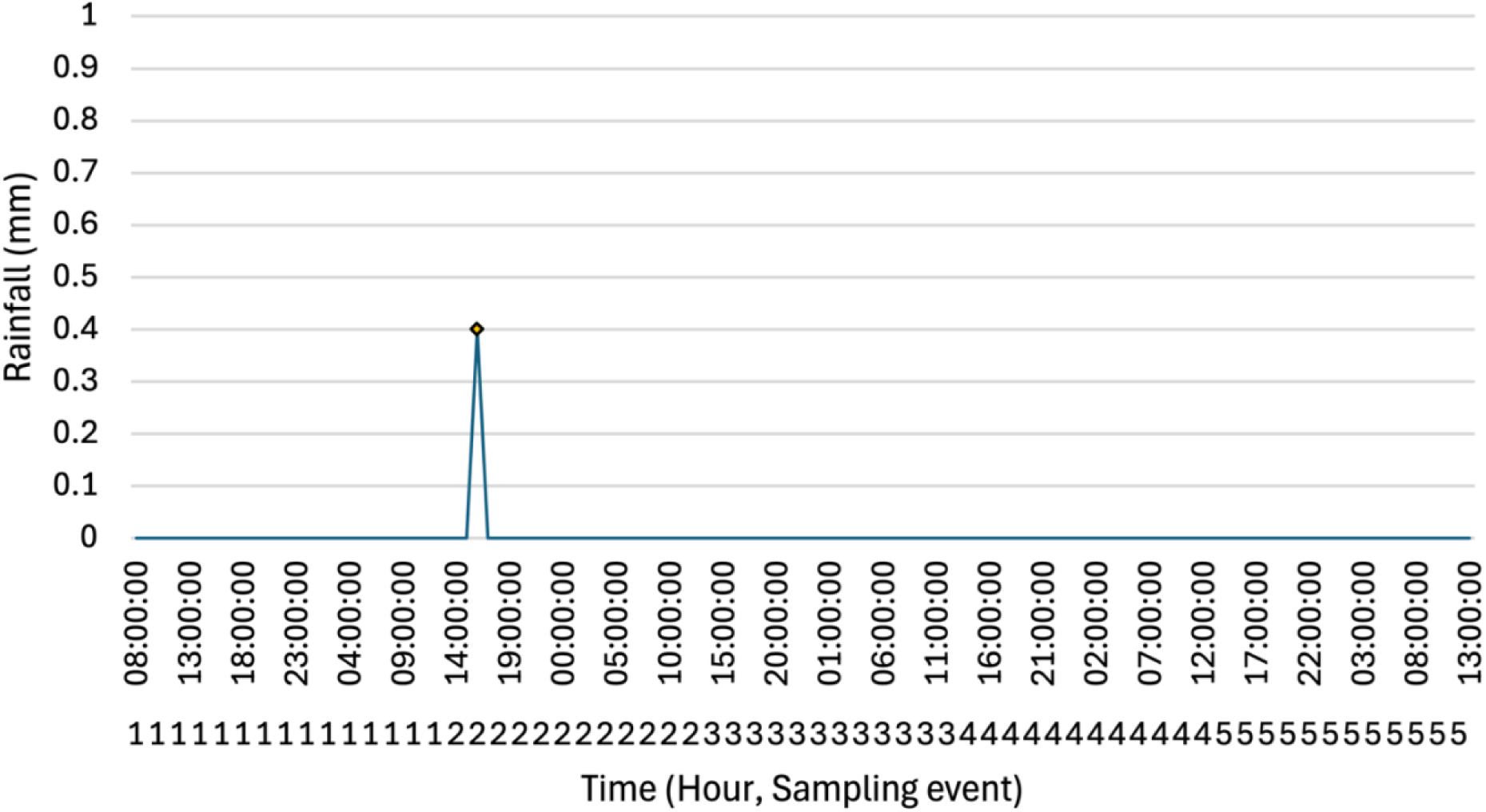
Total rainfall (mm) across five sampling periods of distance experiments 7^th^ – 12^th^ October 2024 Waimate, New Zealand, measured by local weather station set up 1 kilometre distance away from the captive wallaby park EnkleDooVery Korna.

**Figure S7.**
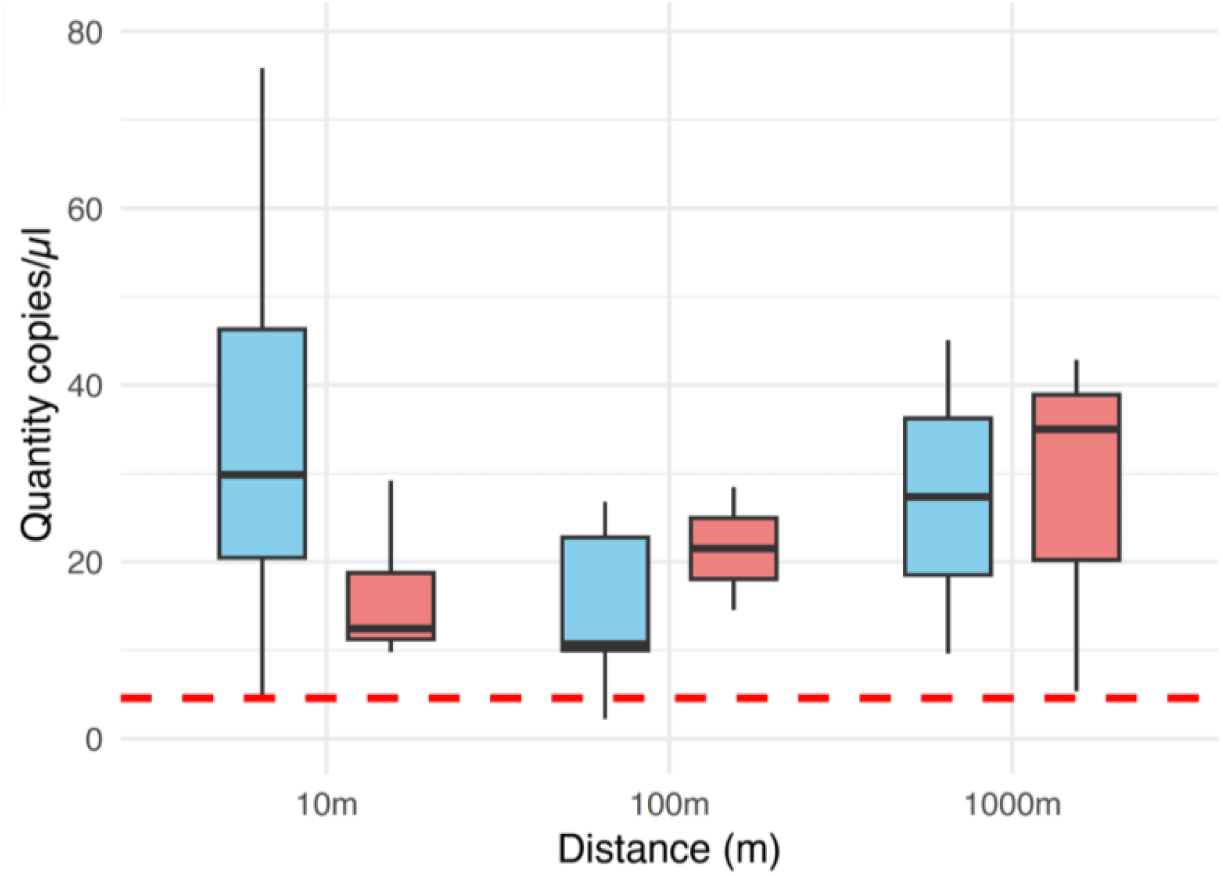
Quantity of target DNA (copies/µL) collected by airborne eDNA sampling using two collection methods: active sampling (blue), and passive sampling (red), at distances ranging from 10-1000 metres from the source of wallaby DNA. The dashed red line indicates the limit of detection (LOD) for this assay; 1 sample fell below this threshold but was retained for analysis. Kruskal-Wallis rank-sum tests show a significant difference in DNA copies/µl of actively collected samples by distance (*p* = 0.0374) but no significant difference in DNA yield for passively collected samples by distance (*p* = 0.6244).

**Figure S8.**
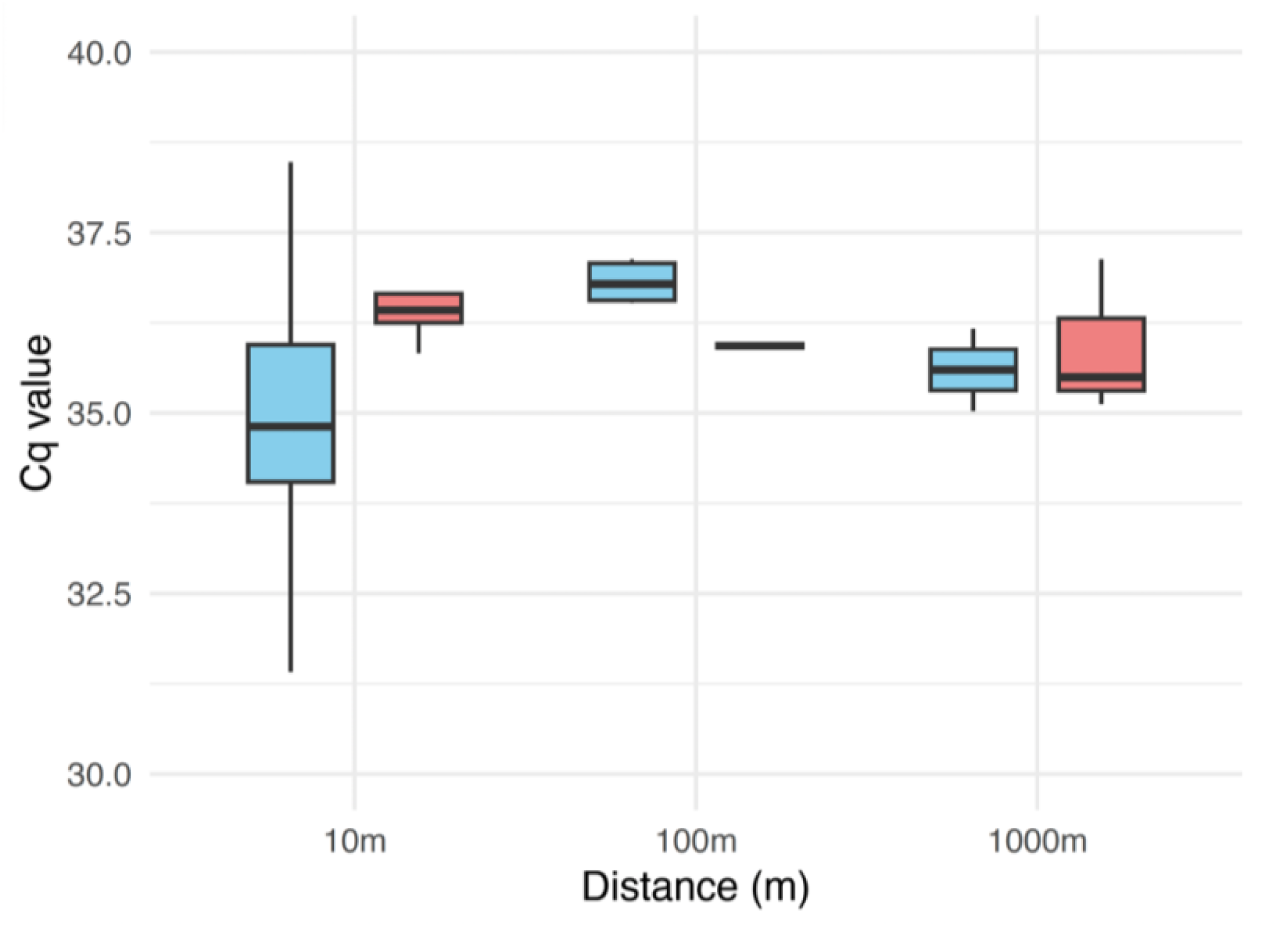
Cycle quantification (Cq) values for airborne eDNA samples using two collection methods: active sampling (blue), and passive sampling (red), at distances ranging from 10-1000 metres from the source of wallaby DNA. Kruskal-Wallis rank-sum tests show a significant difference in Cq value of actively collected samples by distance (*p* = 0.0075) but no significant difference in Cq value for passively collected samples by distance (*p* = 0.5159).

**Table S1.**
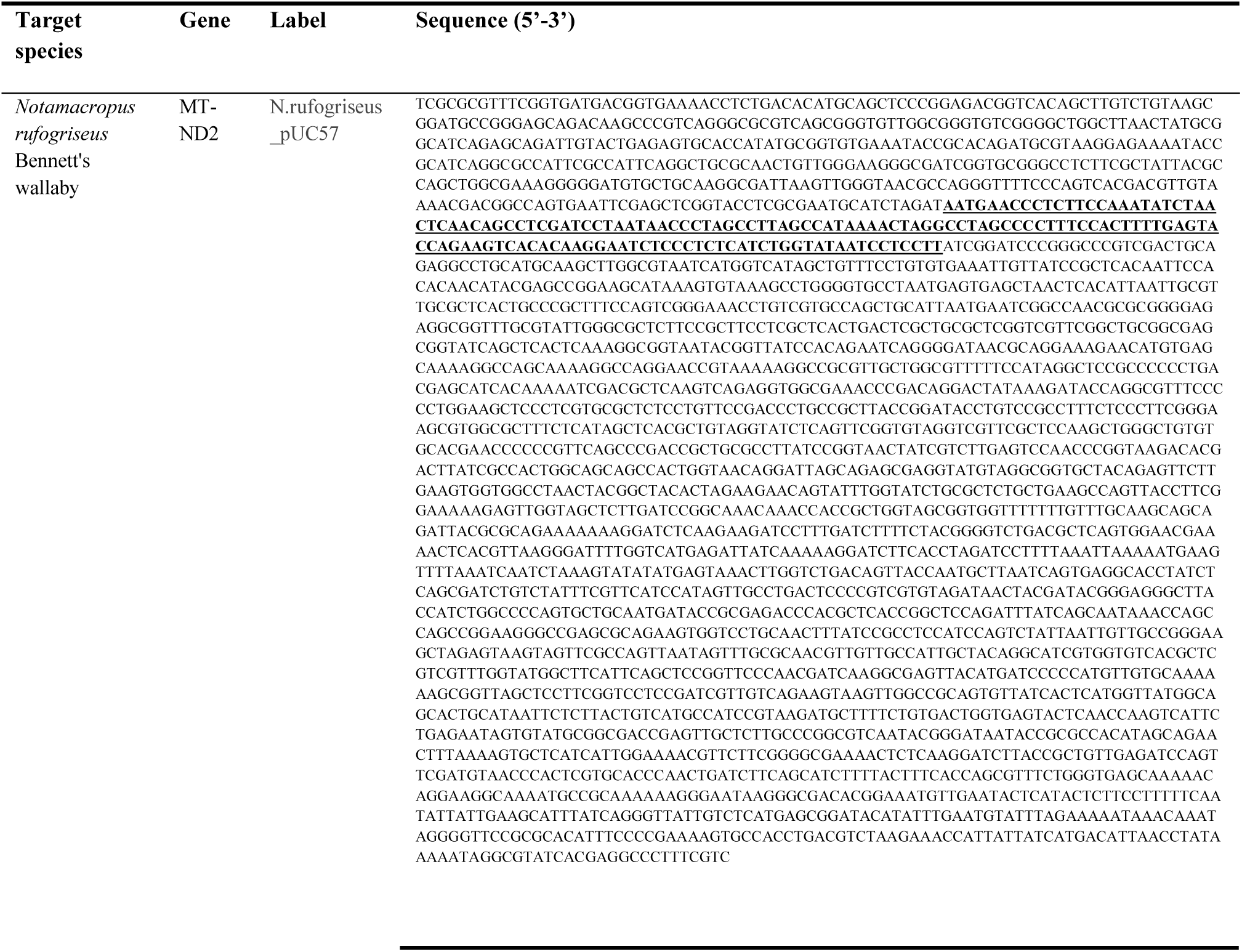
The synthetic oligonucleotide sequence designed to amplify a 146 bp region of the MT-ND2 gene in Bennett’s wallaby *Notamacropus rufogriseus* (underlined and bold) within a pUC57 plasmid, total length 2856 bp.

**Table S2.** Average airflow rates measured using a handheld anemometer for a 12V axial fan with various fibreglass filter configurations: no filter, a G4 grade filter, and an F7 grade filter, both classified according to EN 779:2012 standards. The airflow rates are presented as mean values with standard deviation (SD) in meters per second (m/s), as well as converted to cubic meters per minute (m³/min) and cubic meters per hour (m³/h).

| Filter | Airflow rate m/s ± SD | Airflow rate m³/min ± SD | Airflow rate m³/hour ± SD |
| --- | --- | --- | --- |
| No filter | 5.433 ± 0.294 | 1.216 | 72.98 |
| EN 779:2012 G4 grade | 4.95 ± 0.243 | 1.108 | 66.49 |
| EN 779:2012 F7 grade | 4.1833 ± 0.147 | 0.937 | 56.2 |

**Table S3.**
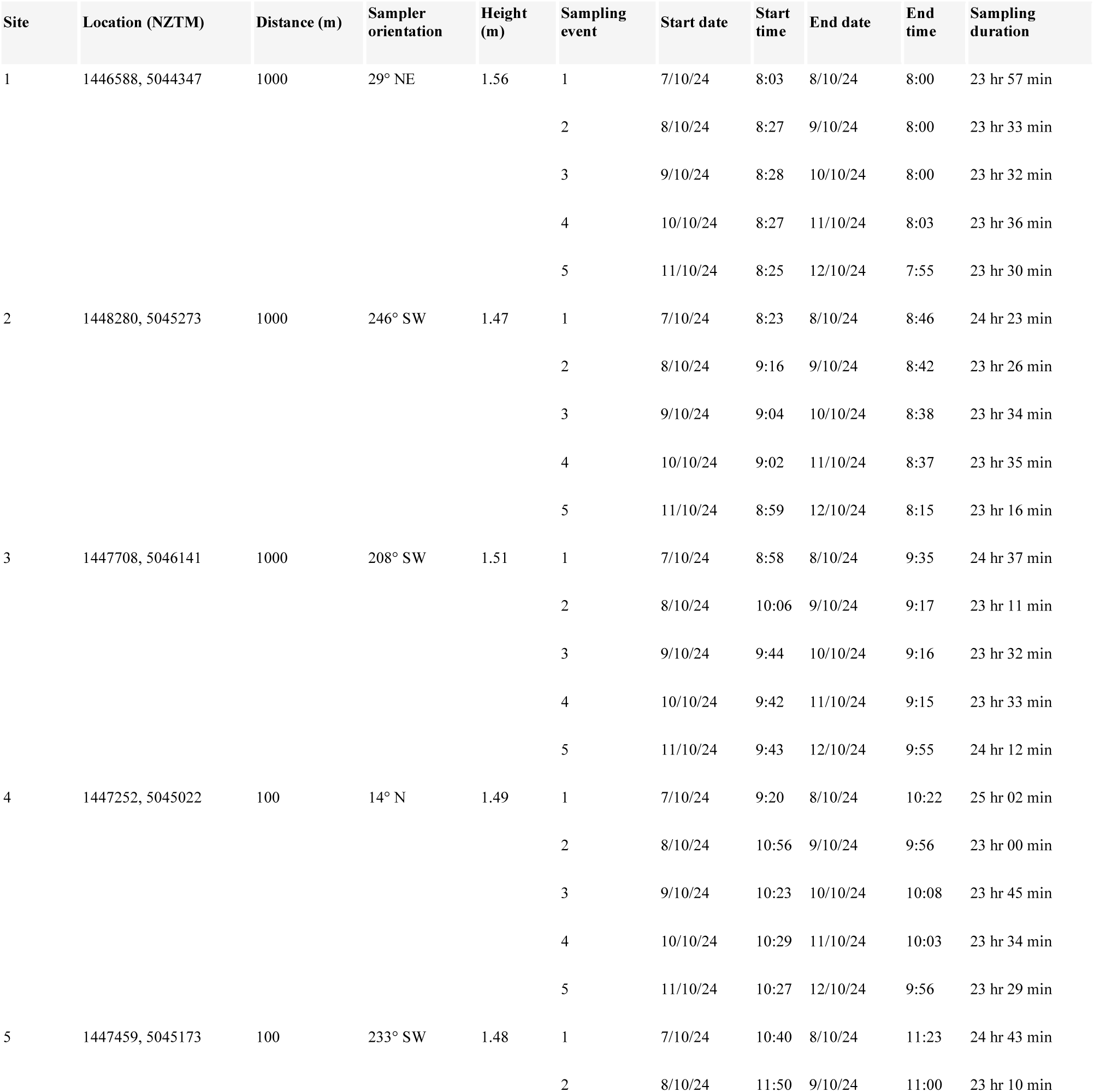
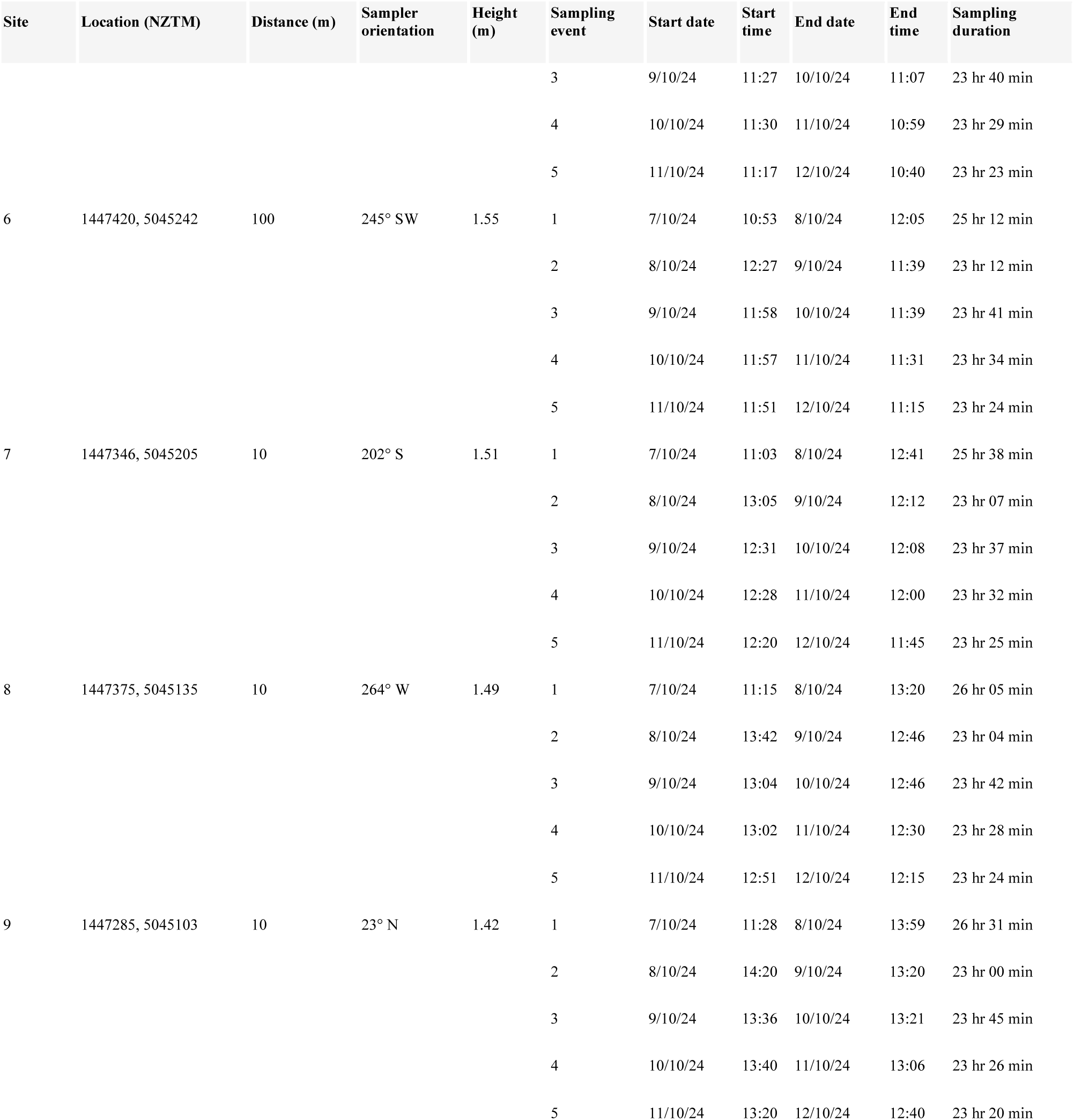
Distance experiment site and sample details. Each row represents an individual airborne eDNA sample, collected over ∼24-hour periods between 7-11 October 2024 in Waimate, South Island, New Zealand. Information includes site ID, location coordinates (NZTM), distance from the source of wallaby DNA at EnkleDooVery Korna tame wallaby park, sampler orientation, height above ground (in metres), sampling event number (1–5), start and end dates and times, and total sampling duration.

**Table S5.** Performance measured using quantitative PCR (qPCR) of the targeted assay at 2 µl reaction volume, using 146 bp region of the MT-ND2 gene in Bennett’s wallaby *Notamacropus rufogriseus* within a pUC57 plasmid (GenScript), total length 2856 bp. The limit of detection (LOD) was determined to be 9.167 copies per 2 µl, while the limit of quantification (LOQ) was 23 copies per 2 µl. The standard curve demonstrated a strong linear relationship with an R² value of 0.991 and a slope of –3.56, corresponding to an amplification efficiency of 90.9%. The lower bound of the 95% confidence interval (Low. 95) was 64.88 copies per 2 µl.

| Parameter | Value |
| --- | --- |
| LOD | 9.167 copies/ 2 µl |
| LOQ | 23 copies/ 2 µl |
| R<sup>2</sup> | 0.991 |
| Slope | -3.56 |
| Low .95 | 64.88 copies/ 2 µl |
| Efficiency | 90.9% |

**Table S6.**
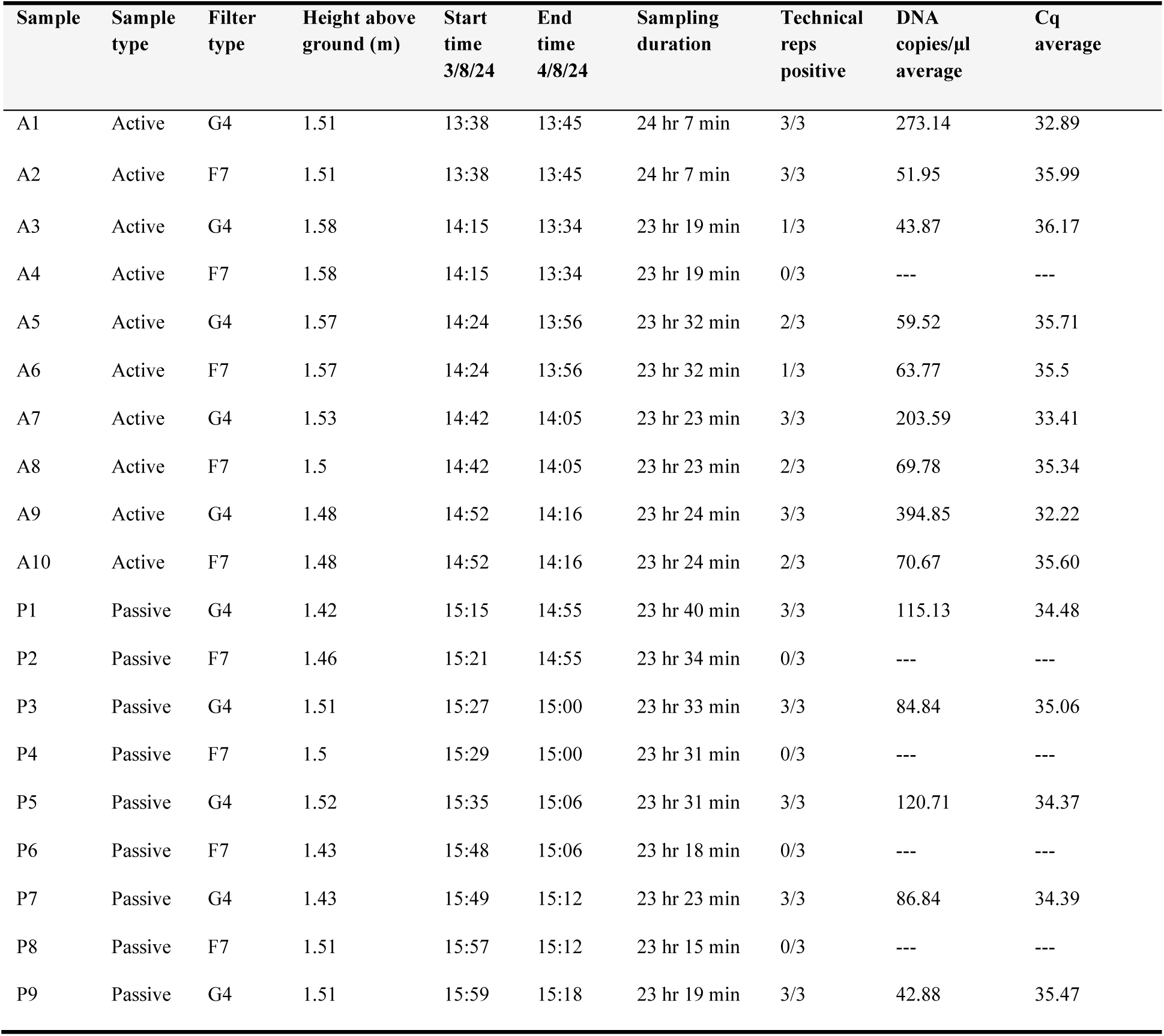
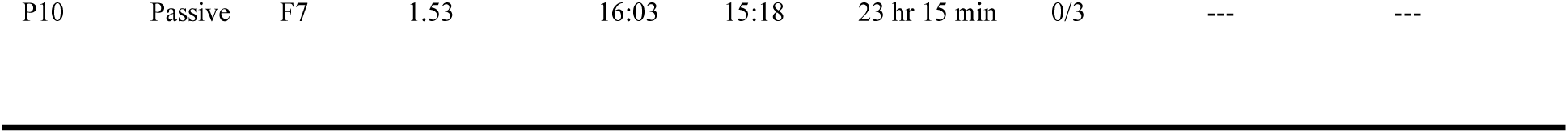
Proof of concept experiment site and sample details. Each row represents an individual airborne eDNA sample, collected over ∼24-hour periods between 3 August and 4 August 2024 in Waimate, South Island, New Zealand. Missing values (e.g., for negative samples) are indicated by dashes.

**Table S7.**
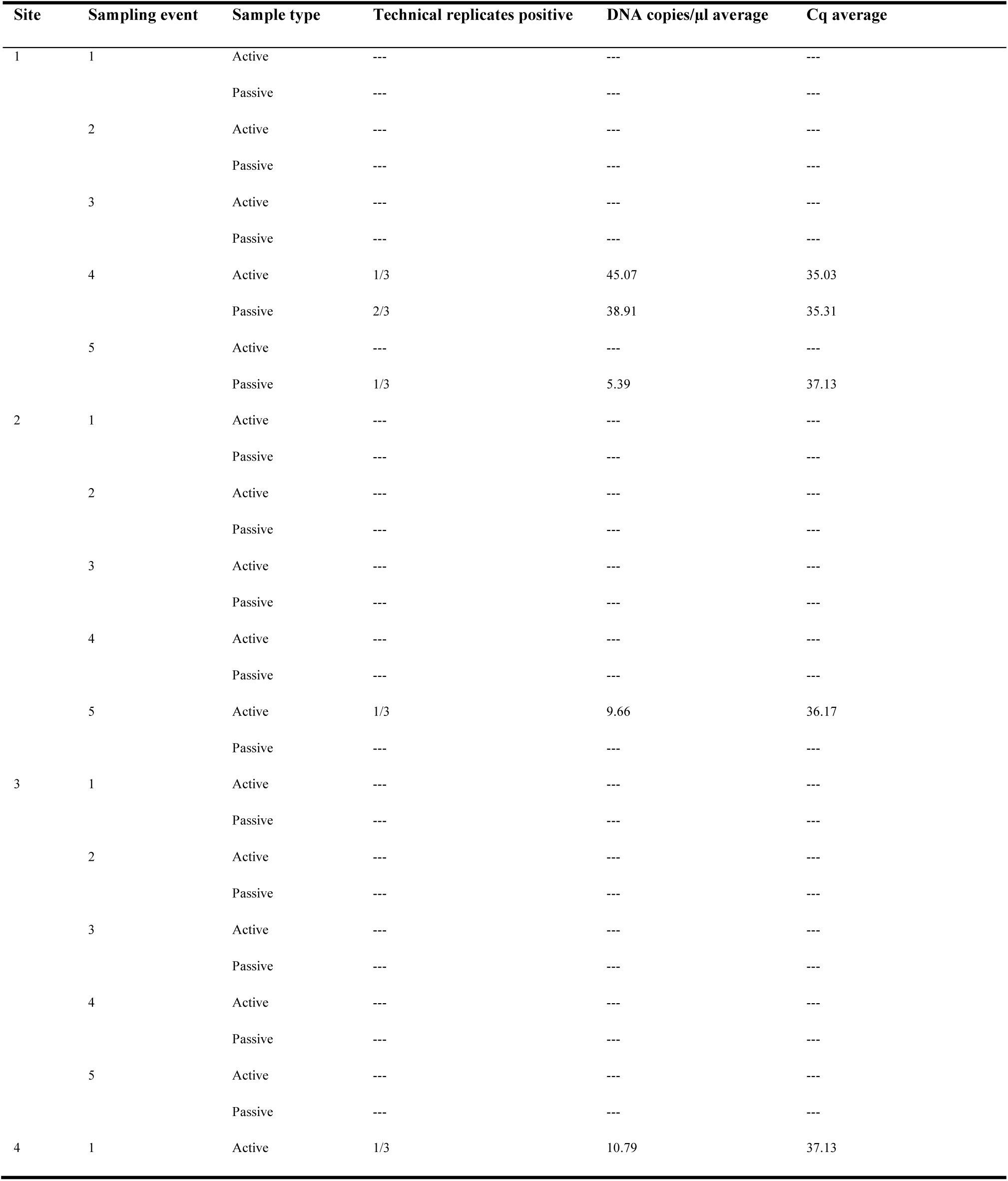
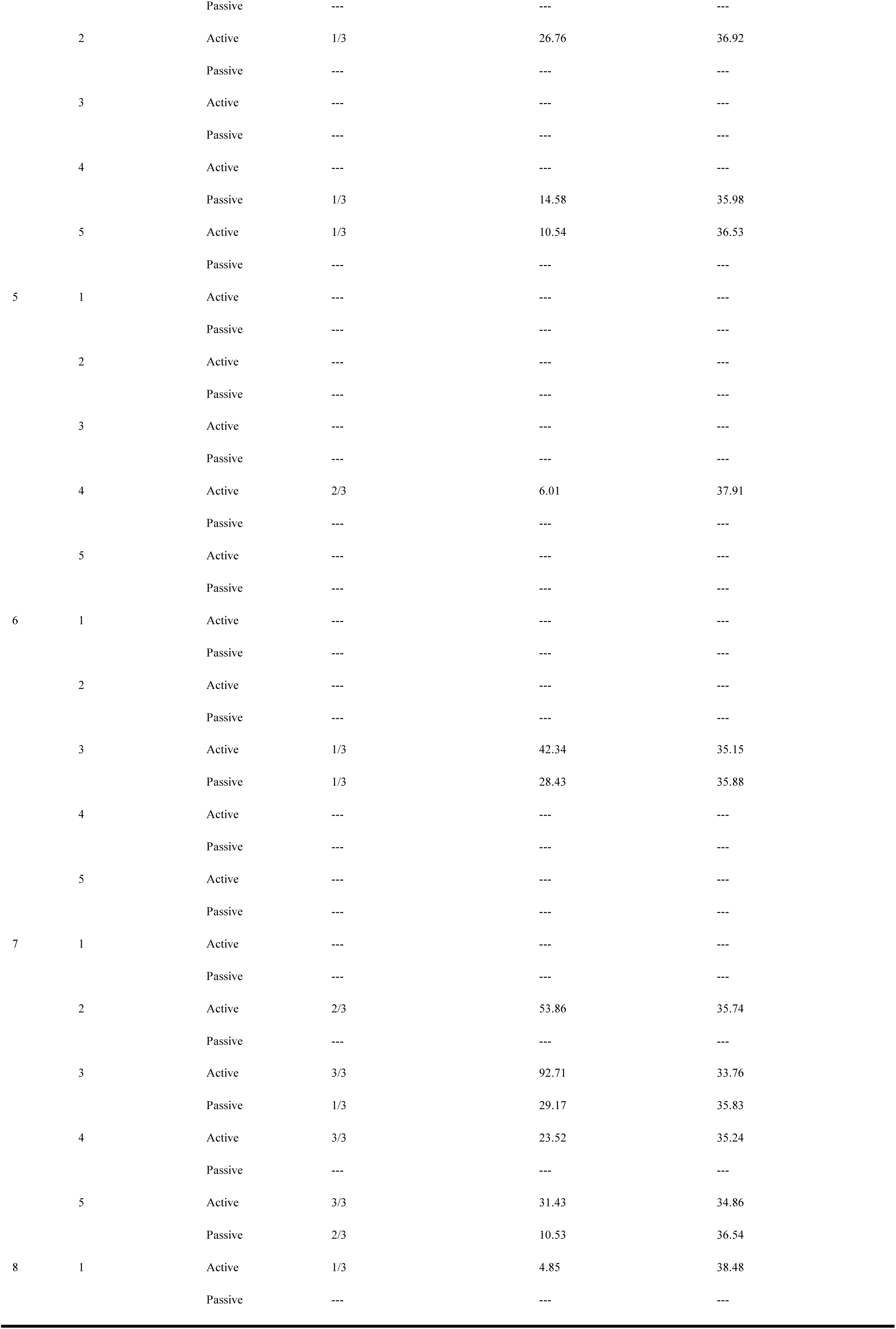
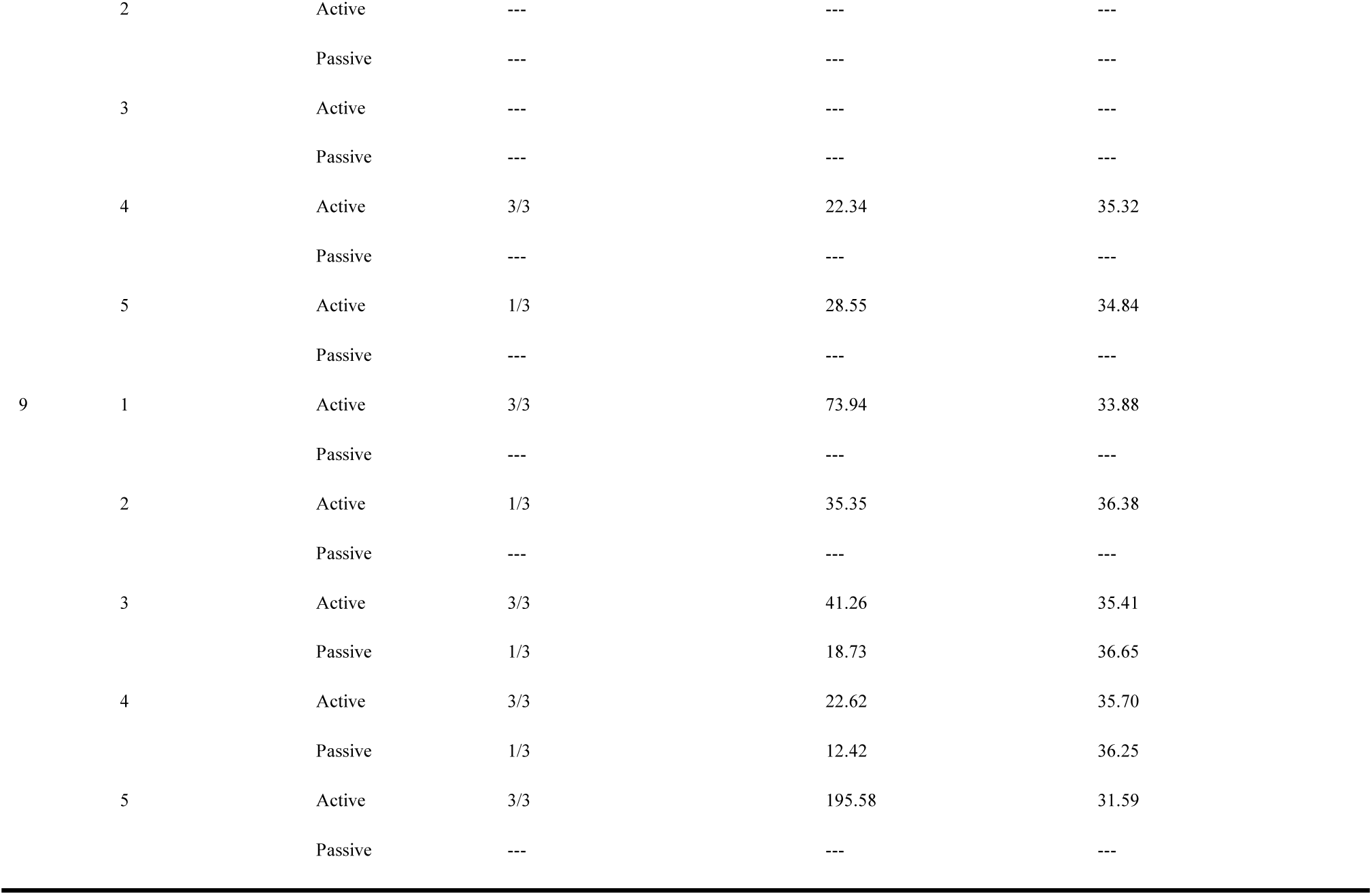
Distance experiment site and sample details. Each row represents an individual airborne eDNA sample, collected over ∼24-hour periods between 7-11 October 2024 in Waimate, South Island, New Zealand. Information includes site ID, sampling event (1-5), sample type (active or passive collection), number of technical replicates testing positive (out of 3), average DNA concentration (copies/µl), and average quantification cycle (Cq) values from qPCR analysis. Missing values for negative samples are indicated by dashes.

**Table S8.** Detection probability (%) and estimated number of replicates required for 95% detection of 1 or 27 wallabies using active (fan-powered) or passive (no fan) airborne eDNA sampling at distances of 10, 100, and 1000 meters from the source of wallaby DNA. Detection probability reflects the likelihood of detecting wallaby DNA at the specified distance from the source using the specified sampling method.

| Sampling Type | Number of wallabies | Distance (m) | Detection Probability (%) | Number of Replicates for 95% Detection |
| --- | --- | --- | --- | --- |
| Active | 27 | 10 | 64.4 | 3 |
| Active | 27 | 100 | 15.6 | 18 |
| Active | 27 | 1000 | 4.4 | 68 |
| Passive | 27 | 10 | 11.1 | 26 |
| Passive | 27 | 100 | 4.44 | 66 |
| Passive | 27 | 1000 | 6.67 | 44 |
| Active | 1 | 10 | 3.75 | 79 |
| Active | 1 | 100 | 0.63 | 477 |
| Active | 1 | 1000 | 0.17 | 1781 |
| Passive | 1 | 10 | 0.43 | 688 |
| Passive | 1 | 100 | 0.17 | 1781 |
| Passive | 1 | 1000 | 0.26 | 1172 |

**Table S9.** Results of the best-performing generalised linear model (GLM) predicting detection probability for airborne eDNA samples. Detection outcomes (positive = 1, negative = 0) were modelled using GLMs with a binomial distribution; fixed effects included distance, environmental variables, and sampler type (active vs. passive). Best-fitting models were selected using backward model selection, based on the lowest Akaike Information Criterion corrected for small sample size (AICc). Models that are considered significant (bolded) have p < 0.05 and 95% bootstrap confidence intervals (CI) excluding zero.

| Fixed effects | Category | Lowest AICc score | Estimate | Pr (>\|z\|) | 95% CI |
| --- | --- | --- | --- | --- | --- |
| Distance | Active samples | 52.63 | -1.34 | **0.0047** | -1826.44, -0.35 |
| Site direction sin |  |  | 1.407 | 0.0397 | -4.18, 368.31 |
| Temperature | Passive samples | 38.97 | -0.825 | 0.0570 | -509.76, 1519.19 |
| Site direction sin |  |  | 1.921 | 0.0275 | -2.01, 9.601e+14 |
| Distance | All samples | 90.88 | -1.072 | **0.0023** | -3.20, -0.34 |
| Sampler type Passive |  |  | -1.619 | **0.0062** | -4.20, -0.46 |
| Temperature |  |  | -0.578 | 0.040 | -1.20, 1.13 |
| Site direction sin |  |  | 1.595 | 0.0023 | -13.08, 12.18 |

**Table S10.** Results of the best-performing Bayesian logistic regression models predicting detection probability for airborne eDNA samples. The model included fixed effects for distance and environmental variables. The model was run with four Markov Chain Monte Carlo (MCMC) chains, resulting in 12,000 post-warmup samples. Convergence was confirmed via R-hat values (∼1.00). Predictors were significant (bolded) when their 95% confidence interval (CI) did not include zero.

| Fixed effects | Category | Estimate (log-odds) | 95% CI |
| --- | --- | --- | --- |
| Distance | Active samples | **-1.17** | **-2.17, -0.15** |
| Temperature |  | -0.43 | -1.14, 0.63 |
| Humidity |  | -0.11 | -1.17, 0.95 |
| Wind speed |  | -0.38 | -1.36, 0.65 |
| Wind direction sin |  | -0.03 | -1.83, 1.78 |
| Wind direction cos |  | -0.06 | -1.93, 1.82 |
| Site direction sin |  | 0.65 | -0.89, 2.11 |
| Site direction cos |  | 0.44 | -0.80, 1.71 |
| Distance | Passive samples | -0.43 | -1.50, 0.59 |
| Temperature |  | -0.66 | -1.80, 0.61 |
| Humidity |  | -0.44 | -1.82, 0.93 |
| Wind speed |  | 0.32 | -0.85, 1.58 |
| Wind direction sin |  | -0.14 | -1.99, 1.73 |
| Wind direction cos |  | -0.09 | -2.02, 1.84 |
| Site direction sin |  | 1.01 | -0.60, 2.58 |
| Site direction cos |  | 0.33 | -0.97, 1.63 |

**Table S11.**
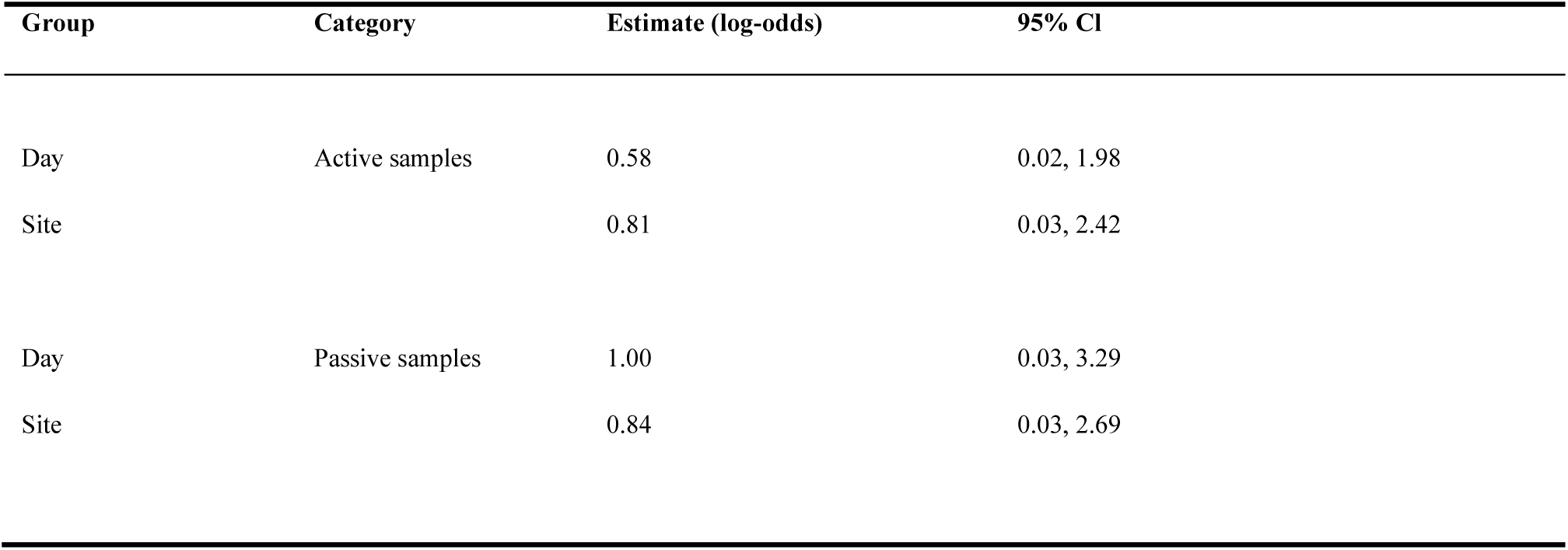
Results of Bayesian logistic regression model random intercepts for sampling site and sampling day to account for spatial and temporal variation. Posterior standard deviations (SD) of the random intercepts are presented with their associated 95% confidence intervals (CI). Estimates are based on 12,000 post-warmup samples from four MCMC chains.

| Group | Category | Estimate (log-odds) | 95% CI |
| --- | --- | --- | --- |
| Day | Active samples | 0.58 | 0.02, 1.98 |
| Site |  | 0.81 | 0.03, 2.42 |
| Day | Passive samples | 1.00 | 0.03, 3.29 |
| Site |  | 0.84 | 0.03, 2.69 |

